# Proteins Derived From MRL/MpJ Tendon Provisional Extracellular Matrix and Secretome Promote Pro-Regenerative Tenocyte Behavior

**DOI:** 10.1101/2024.07.08.602500

**Authors:** Jason C. Marvin, Ethan J. Liu, Hsin Huei Chen, David A. Shiovitz, Nelly Andarawis-Puri

## Abstract

Tendinopathies are prevalent musculoskeletal conditions that have no effective therapies to attenuate scar formation. In contrast to other adult mammals, the tendons of Murphy Roths Large (MRL/MpJ) mice possess a superior healing capacity following acute and overuse injuries. Here, we hypothesized that the application of biological cues derived from the local MRL/MpJ tendon environment would direct otherwise scar-mediated tenocytes towards a pro-regenerative MRL/MpJ-like phenotype. We identified soluble factors enriched in the secretome of MRL/MpJ tenocytes using bioreactor systems and quantitative proteomics. We then demonstrated that the combined administration of structural and soluble constituents isolated from decellularized MRL/MpJ tendon provisional ECM (dPECM) and the secretome stimulate scar-mediated rodent tenocytes towards enhanced mechanosensitivity, proliferation, intercellular communication, and ECM deposition associated with MRL/MpJ cell behavior. Our findings highlight key biological mechanisms that drive MRL/MpJ tenocyte activity and their interspecies utility to be harnessed for therapeutic strategies that promote pro-regenerative healing outcomes.

**Teaser:** Proteins enriched in a super-healer mouse strain elicit interspecies utility in promoting pro-regenerative tenocyte behavior.

## Introduction

Tendinopathies are debilitating disorders that account for over a third of all musculoskeletal consultations (1). In adult mammals, tendon ruptures heal poorly by forming fibrovascular and mechanically inferior scar tissue (‘scar-mediated healing’) that is later remodeled ineffectively by resident tendon cells, or tenocytes. This scar-mediated healing can result in long-term pain, disability, and functional deficits that contribute to a reduced quality of life. Surgical repair remains the gold standard for the clinical treatment of tendon ruptures wherein the torn ends are reattached to restore load transmission and the original tissue functionality. However, these procedures do not address the underlying pathology and suffer from post-operative re-tear rates as high as 94% (2–4). The use of biologic therapies, such as autologous platelet-rich plasma (PRP) injections, to promote tendon healing remains contentious given variable reports of their clinical efficacy in attenuating scar-mediated healing (5,6), which is attributed to a lack of standardization in their manufacturing, quality control, and administration. Clearly, a better understanding the biological mechanisms underpinning tendon regeneration is critical to development of more effective therapeutics.

The Murphy Roths Large (MRL/MpJ) mouse strain harbors a natural regenerative capacity in several tissues and organs (7,8), including tendon (9–15). We have previously shown that MRL/MpJ tendons exhibit vastly improved structural and mechanical recovery as compared to canonical healing C57BL/6 (B6) tendons following acute laceration/full-thickness hole punch or overuse injuries (9,12,13), thereby establishing the promise of the MRL/MpJ mouse strain for expanding our mechanistic understanding of scarless tendon healing. Using *ex vivo* organ culture to eliminate any systemic contributions to healing, we previously found that hole-punched MRL/MpJ tendons showed significantly increased bulk mechanical properties compared to those of B6 tendons at 4 weeks post-injury (wpi). These findings suggest that the local tendon environment consisting of resident tenocytes and the extracellular matrix (ECM) primarily drive MRL/MpJ scarless tendon healing (11,12), which raised the question of whether the MRL/MpJ tendon ECM can ameliorate otherwise scar-mediated healing outcomes. Subsequent *in vivo* assessment of the therapeutic utility of a PEG-4MAL hydrogel incorporated with decellularized MRL/MpJ tendon provisional ECM (dPECM), a temporary bioactive ECM produced during the proliferative phase of healing, improved B6 tendon alignment, cell elongation, and stiffness compared to untreated tendons by 6 weeks post-implantation (16). Moreover, in that study, we identified a subset of proteins that are uniquely enriched in the MRL/MpJ tendon dPECM that may contribute to these improved healing outcomes. However, native tissue mechanical properties were not fully recovered despite the enhanced structure achieved using these dPECM constructs. While decellularization of tissue samples for dPECM collection is necessary to prevent immunogenicity, this technique removes resident tenocytes and most of their direct contributions within the tendon. These findings suggest that supplementation of tenocyte-secreted soluble factors, otherwise known as the secretome, may augment the therapeutic potency of the MRL/MpJ tendon dPECM.

This study aims to (*i*) comparatively characterize B6 and MRL/MpJ tenocyte behavior to establish robust cellular metrics that delineate scar-mediated and scarless tendon healing, (*ii*) identify soluble factors enriched in the MRL/MpJ tenocyte secretome, and (*iii*) determine whether the administration of MRL/MpJ-enriched soluble and ECM proteins together is functionally advantageous in stimulating pro-regenerative MRL/MpJ-like activity compared to either constituent alone. We addressed these goals by adopting an experimental approach integrating primary cell culture, functional *in vitro* assays, and *in vivo* histological analyses that capture physiological aspects of acute tendon healing and quantitative proteomics to profile the secretome. We hypothesized that harnessing biological cues derived from the local MRL/MpJ tendon environment through recombinant protein design would recapitulate the beneficial effects of administering these MRL/MpJ constituents *in vitro.* Our findings shed insight into the cellular processes and candidate protein regulators that drive effective mammalian tendon healing across rodent species, establishing a novel biologics-based therapeutic strategy that also provides a promising platform for future mechanistic interrogation.

## Results

### MRL/MpJ tenocytes demonstrate greater intercellular communication, cytoskeletal organization, mechanosensitivity, and ECM deposition than B6 tenocytes

We have previously shown that culturing B6 tenocytes on MRL/MpJ dPECM-coated 2D substrates attained a dendritic morphology consisting of axon-like protrusions and decreased circularity(12). To ascertain whether these protrusions are reflective of MRL/MpJ tenocyte behavior, we analyzed the single-cell morphology of B6 and MRL/MpJ tenocytes cultured on collagen I-coated substrates using phalloidin staining (**Fig. 1A and B**). Compared with B6 tenocytes, a significantly greater proportion of MRL/MpJ tenocytes formed actin-rich protrusions (70.3% compared to 88.9% of all cells, *P* = 0.034; **Fig. 1C**). There were no differences in the number of protrusions formed on a per cell basis between B6 and MRL/MpJ tenocytes (data not shown), indicating that the significantly greater total protrusion length (total and normalized to cell spreading area; *P* < 0.0001 and 0.0029, respectively) observed in MRL/MpJ tenocytes (**Fig. 1C**) is due to longer cytoskeletal extensions. Importantly, these results confirm that our prior observations are characteristic of MRL/MpJ tenocyte morphology that may mediate long-distance cell-cell contacts and could be suggestive of increased mechanosensitivity. Additionally, MRL/MpJ tenocytes also showed significantly lower cell circularity (*P* = 0.0001) and greater spreading area (*P* = 0.03) and perimeter (*P* < 0.0001) (**Fig. 1D**), which corroborates our prior morphological observations in B6 tenocytes cultured on MRL/MpJ dPECM also as characteristic of MRL/MpJ cell behavior. We next investigated whether these intrinsic differences in the actin cytoskeleton serve as a physical mediator to enable MRL/MpJ tenocytes to better engage with changes to their surrounding microenvironment.

**Fig. 1.**
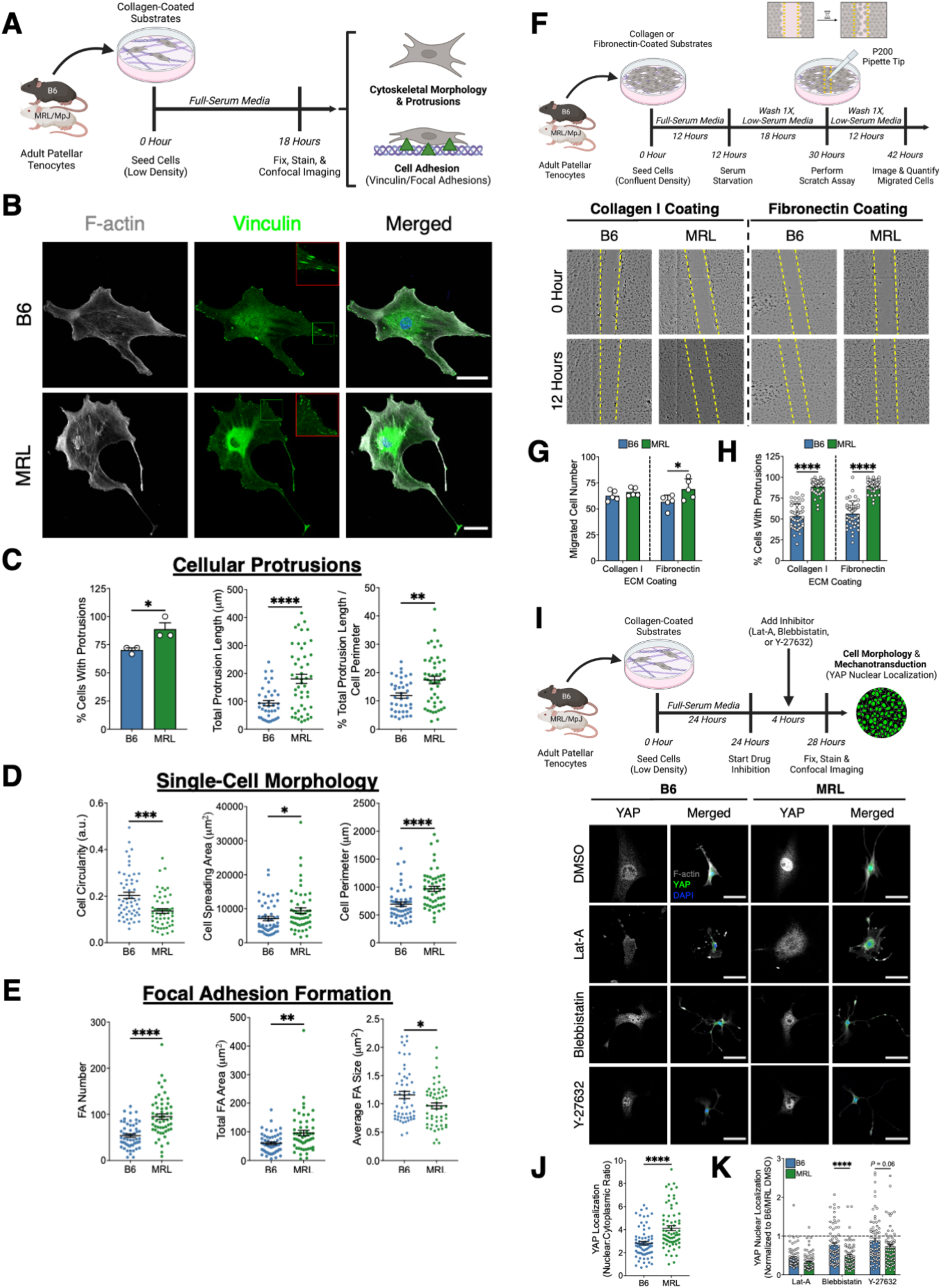
MRL/MpJ tenocytes demonstrate enhanced mechanosensitivity and cell migration and form more actin-rich, axon-like cellular protrusions compared to B6 tenocytes. (**A**) Experimental design to assess characterize morphology and focal adhesion formation. (**B to E**) IF staining of F-actin (gray) and vinculin (green) of B6 and MRL/MpJ tenocytes cultured on collagen I-coated substrates (scale bar, 100 μm; 2X-magnified insets in the top-right corner of representative vinculin images). MRL/MpJ tenocytes formed more extensive cellular protrusions (C) that were accompanied by greater cell spreading and perimeter (D) and assembly of nascent focal adhesions compared to B6 tenocytes (E). (**F to H**) Experimental design to quantify tenocyte migration via wound scratch assays. Wound scratch analysis (F) showed that a significantly greater number of MRL/MpJ tenocytes than B6 tenocytes migrated on fibronectin-coated substrates after 12 hours post-scratch (G) despite comparable cell protrusion characteristics between cells cultured on collagen I and fibronectin (H). (**I**) Experimental design to assess YAP-mediated mechanosignaling. IF staining of phalloidin (grey) and YAP (magnified 3X in gray; green in merged representative images) of B6 and MRL/MpJ tenocytes cultured on collagen I-coated substrates (scale bar, 100 μm) with pharmacological inhibitors for actin polymerization (Lat-A), myosin II (blebbistatin), and ROCK (Y-27632). DMSO-treated cells served as a control. (**J**) At baseline, MRL/MpJ tenocytes displayed significantly higher YAP nuclear localization than B6 tenocytes. (**K**) Relative nuclear translocation of YAP (normalized to respective DMSO control mean) was significantly reduced in MRL/MpJ tenocytes compared to B6 tenocytes with inhibition of myosin II and ROCK. *\*P* < 0.05, \*\**P* < 0.01, \*\*\**P* < 0.001, \*\*\*\**P* < 0.0001. Significance was determined using a Welch’s t-test when comparing B6 and MRL/MpJ tenocytes, or a two-way ANOVA with post-hoc Bonferroni when comparing genotypes for each inhibitor for YAP data. Representative images shown for all staining experiments.

It is well established that the contractile actomyosin machinery is required to drive changes in cell shape, adhesion, and motility that regulate cell signaling and behavior (17). In particular, actin-based cellular protrusions facilitate cell migration and spreading and are closely coupled with the formation of focal adhesions (FAs) such as vinculin. Vinculin regulates the recruitment and disassembly of these FAs that link the actin cytoskeleton to the ECM and enable cells to sense and respond to environmental factors (e.g., activation of growth factor-mediated signaling pathways, mechanical cues) (18–21). Thus, we hypothesized that the protrusion networks that are characteristic of MRL/MpJ tenocytes enhance mechanosensitivity and cell migration. Indeed, MRL/MpJ tenocytes exhibited significantly greater FA number (1.77-fold higher, *P* < 0.0001) and area (1.6-fold higher, *P* = 0.0012) than that of B6 tenocytes when cultured on collagen I-coated substrates (**Fig. 1E**). However, when normalized to cell spreading area, FA area was not statistically different between B6 and MRL/MpJ cells (data not shown; *P* = 0.22), but FA number remained significantly higher in MRL/MpJ tenocytes (*P* = 0.001). Moreover, the average FA size was significantly greater in B6 tenocytes than MRL/MpJ tenocytes (1.16 ± 0.064 μm^2^ compared to 0.96 ± 0.05 μm^2^), indicating that these MRL/MpJ protrusions are stabilized by smaller nascent adhesions (20) (<1 μm diameter). These findings indicate that MRL/MpJ tenocytes undergo rapid and robust turnover of smaller FAs that better stabilize the actin cytoskeleton and its integrin-mediated mechanotransduction compared to B6 tenocytes. Furthermore, 2D wound scratch assays (**Fig. 1F**) showed significantly increased MRL/MpJ tenocyte numbers at the defect site compared to B6 tenocytes on fibronectin-coated substrates (*P* = 0.048; **Fig. 1G**). Cell migration was not different on collagen I-coated substrates (*P* = 0.25) despite comparable protrusion networks on both collagen I and fibronectin (**Fig. 1H**), indicating that changes to other cell-ECM and/or cell-cell interactions during an injury response likely play a more important role in modulating 2D tenocyte migration.

Cytoskeletal tension-induced activation of Hippo pathway effectors and transcriptional coactivators, Yes-associated protein (YAP) and WW domain-containing transcription regulator protein 1 (TAZ), regulates cell mechanosensing and adhesion-mediated signaling (22–25). To ascertain whether our observations of increased stress fiber, FA, and cellular protrusion formation in MRL/MpJ tenocytes is indicative of a YAP-dependent mechanosignaling program, we next evaluated the effects of inhibiting actin stabilization and ROCK-mediated cell contractility on YAP nuclear translocation. We cultured B6 and MRL/MpJ tenocytes at low density on collagen I-coated substrates with and without Latrunculin A (inhibitor of actin polymerization), blebbistatin (inhibitor of myosin II, a downstream effector of ROCK), and Y-27632 (inhibitor of ROCK) for 4 hours and stained for YAP (**Fig. 1I**). As expected, YAP nuclear translocation was significantly greater in MRL/MpJ tenocytes than B6 tenocytes (1.46-fold higher, *P* < 0.0001) treated with the DMSO vehicle control (**Fig. 1J**). No correlations were found between YAP nuclear translocation and single-cell morphological parameters. To account for baseline differences in YAP nuclear translocation between B6 and MRL/MpJ tenocytes, we subsequently normalized data relative to the DMSO control and compared the relative changes in YAP nuclear translocation with pharmacological inhibition between mouse strains (**Fig. 1K**). Unsurprisingly, disruption of the actin cytoskeleton with Latrunculin A treatment resulted in YAP being predominantly localized in the cytoplasm with no difference in relative YAP nuclear translocation between B6 and MRL/MpJ tenocytes (*P* = 0.22), indicating that actin polymerization is required for nuclear transport of YAP. Inhibition of myosin II with blebbistatin treatment significantly reduced relative YAP nuclear translocation in MRL/MpJ tenocytes compared to B6 tenocytes (*P* < 0.0001) with MRL/MpJ tenocytes having greater cytoplasmic retention of YAP. Similarly, inhibition of ROCK with Y-27632 treatment reduced relative YAP nuclear translocation in MRL/MpJ tenocytes compared to B6 tenocytes (*P* = 0.058). These data confirm that MRL/MpJ tenocytes harbor enhanced mechanosignaling dependent on myosin II- and ROCK-mediated contractility to facilitate YAP nuclear translocation.

We next extended our characterization of B6 and MRL/MpJ cell behavior to confluent culture conditions that are more reflective of the dPECM environment during the proliferative phase of acute tendon healing at 7 dpi *in vivo* (11). After 7 days of culture in full-serum media to reach confluency (**Fig. 2A**), cytoskeletal and cell-deposited fibronectin coherency/alignment (**Fig. 2B-D**) were significantly greater in MRL/MpJ tenocytes than B6 tenocytes (both *P* < 0.0001), consistent with early improvements in cellular and ECM organization seen at 7 dpi *in vivo* in MRL/MpJ tendons compared to B6 tendons (11). Remarkably, many MRL/MpJ nuclei also assembled into tightly-packed, linear arrays along fibronectin tracks (yellow dashed line; **Fig. 2C**) that closely resembled physiological tendon organization despite the absence of structural or mechanical cues during culture. MRL/MpJ tenocytes also exhibited significantly greater ECM deposition (1.46- and 1.29-fold higher for fibronectin and collagen I, respectively; *P* < 0.0001 for both) and thickness (*P* = 0.0021 for fibronectin), cell number (1.50-fold higher, *P* < 0.0001), and expression of connexin-43 (Cx-43) gap junctions (*P* = 0.0089) compared to B6 tenocytes (**Fig. 2E and F**). Moreover, the increasingly anisotropic and fibrillar fibronectin network generated by MRL/MpJ tenocytes is consistent with ROCK- and mechanosignaling-mediated (e.g., ITGB1, FAK) pathways being known to regulate fibronectin assembly (26–29). Visually, fibronectin and Cx-43 gap junctions were more uniformly distributed throughout neighboring MRL/MpJ tenocytes, whereas these were limited to the periphery of B6 tenocytes suggestive of disrupted cell-ECM adhesion (yellow arrowheads, **Fig. 2C**). However, it is not clear whether increased Cx-43 gap junction formation is characteristic of MRL/MpJ tenocyte behavior or driven by changes in cell density. To overcome this limitation, we next cultured cells for 3 days in full-serum media to yield comparable densities. Cx-43 expression remained significantly higher in MRL/MpJ tenocytes than B6 cells (1.32-fold higher, *P* = 0.024) and predominantly localized at the tips of actin-based cellular protrusions at cell-cell contact sites (yellow arrowheads, **Fig. 2G**). There were no differences in αSMA expression after 3 (*P* = 0.6) and 7 (2.2% lower, *P* = 0.14) days of culture (**Fig. 2H**), indicating that increased cytoskeletal organization of MRL/MpJ tenocytes at confluency is not primarily mediated by myofibroblast activity. In short, our data demonstrate that enhanced cytoskeletal and ECM alignment, ECM deposition, and intercellular communication are fundamental to MRL/MpJ tenocyte behavior and likely contribute to improved structural healing outcomes.

**Fig. 2.**
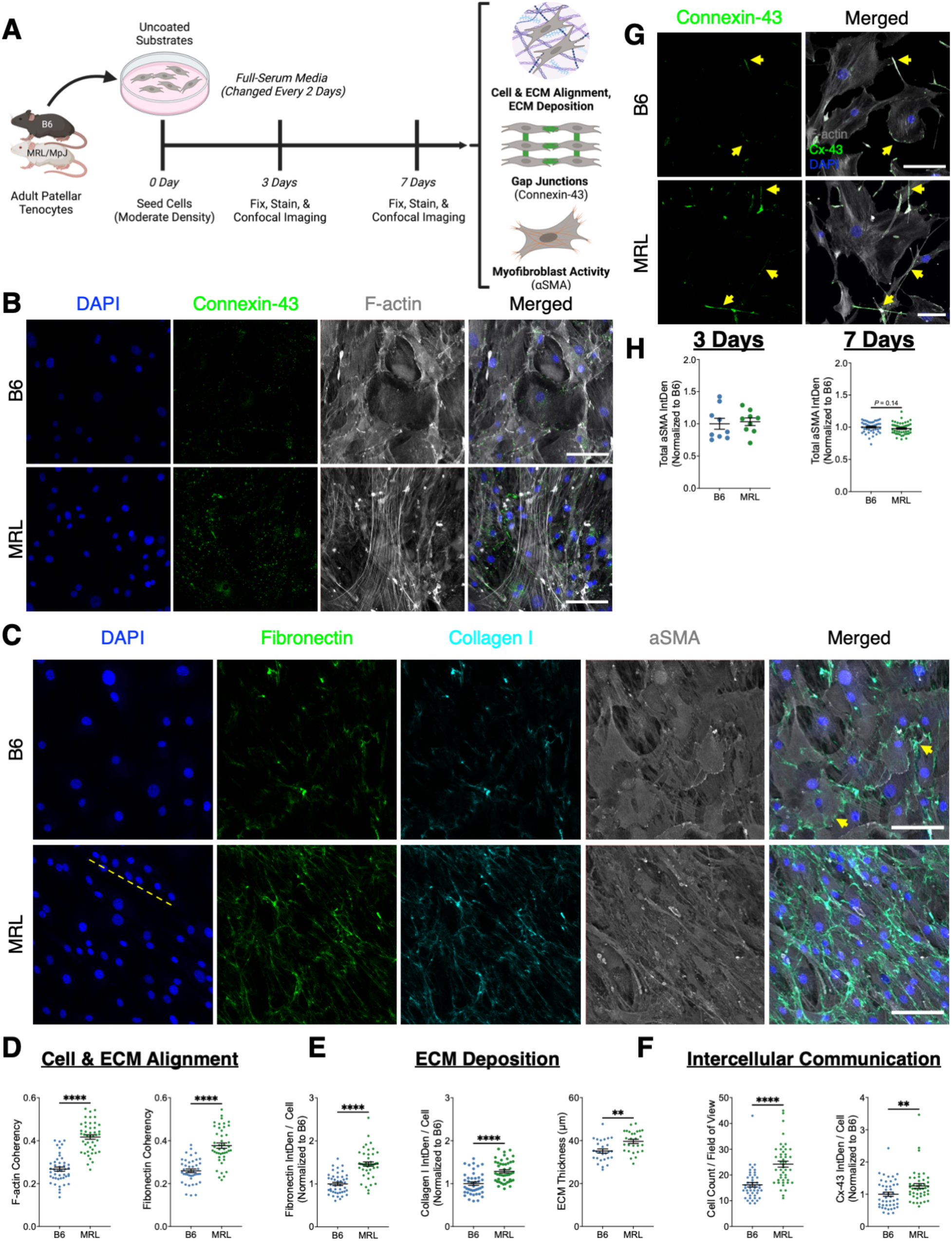
MRL/MpJ tenocytes promote cytoskeletal and cellular alignment, ECM deposition, and intercellular communication consisted with effective tendon healing outcomes *in vivo*. (**A**) Experimental design to assess confluent tenocyte behavior after 7 days of culture in full-serum media. (**B to D**) IF staining of (B) connexin-43 (Cx-43, green) and F-actin (grey) in addition to (C) fibronectin (green), collagen I (cyan), and αSMA (gray) in confluent B6 and MRL/MpJ tenocytes (scale bar, 100 μm). MRL/MpJ tenocytes displayed greater cytoskeletal and matrix anisotropy, ECM fibrillogenesis, and arrangement of nuclei in linear arrays (indicated by the dashed yellow line) compared to B6 tenocytes. (**D to F**) MRL/MpJ tenocytes exhibited significantly greater F-actin and fibronectin coherency/alignment (D), fibronectin and collagen I deposition (E), and cell number and Cx-43 levels (F) than B6 tenocytes. (**G**) IF staining of Cx-43 (green) and F-actin (grey) at moderate cell densities after 3 days of culture revealed that gap junctions localize along the length and tips of actin-rich cellular protrusions (indicated by yellow arrowheads) to facilitate intercellular communication. (**H**) αSMA expression remained comparable between B6 and MRL/MpJ tenocytes after 3 and 7 days of culture suggesting that increased myofibroblast activation does not drive improved structural organization *in vitro*. *\*P* < 0.05, \*\**P* < 0.01, \*\*\**P* < 0.001, \*\*\*\**P* < 0.0001. Significance was determined using a Welch’s t-test when comparing coherency, ECM thickness, or cell number data. Integrated density data on a per cell basis further normalized to the B6 mean was compared using a non-parametric Mann-Whitney *U* test. Representative images shown for all staining experiments.

Given these cell-intrinsic differences in B6 and MRL/MpJ tenocyte morphology and behavior, we next used quantitative proteomics to identify growth factors and cytokines enriched in the MRL/MpJ tenocyte secretome.

### The MRL/MpJ tenocyte secretome contains distinct biological signatures to those of B6 tenocytes regardless of loading modality

We first comprehensively characterized soluble proteins in B6 and MRL/MpJ tenocyte secretomes by profiling concentrated static and dynamic (i.e., cells conditioned under cyclic stretch using the CellScale MCFX bioreactor system) conditioned media samples for up to 640 and 200 protein targets, respectively, using Quantibody Mouse Cytokine Arrays (**Fig. 3A**). Due to sample volume constraints using the MCFX bioreactor compared to traditional culture flasks, quantitative proteomics of the dynamic secretome was restricted to 200 targets. There were no differences in protein concentration (**Fig. S1A**) or cell number at the time of conditioned media collection (data not shown) between B6 and MRL/MpJ secretome samples. In the static secretome (**Fig. 3B**), we identified 336 cytokines of which 252 (75% of detected proteins) were differentially expressed proteins (DEPs) between B6 (50.9% of detected proteins, 118 enriched in B6, 53 only detected in B6) and MRL/MpJ tenocytes (24.1% of detected proteins, 74 enriched in MRL/MpJ, 7 only detected in MRL/MpJ). In the dynamic secretome (**Fig. 3B**), we identified 56 cytokines of which 43 (76.8% of detected proteins) were differentially expressed between B6 (44.6% of detected proteins, 21 enriched in B6, 4 only detected in B6) and MRL/MpJ tenocytes (32.1% of detected proteins, 17 enriched in MRL/MpJ, and 1 only detected in MRL/MpJ). Of note, interleukin-3 (IL-3) was only detected in the B6 secretomes, tumor necrosis factor alpha (TNF-α) was only detected in the dynamic secretomes, and TNF receptor 2 (TNF RII), thymic stromal lymphopoietin (TLSP), and nephrilysin were only detected in the static secretomes. We applied multivariate analyses using principal components analysis (PCA; **Fig. 3C**) and found that B6 and MRL/MpJ static secretomes formed separate clusters along PC1 (34.4% of total variance). PCA revealed minor overlap of B6 and MRL/MpJ dynamic secretome samples reflective of higher biological variation. Heatmap visualization corroborated these observations as evidenced by a greater degree of sample clustering in the static secretomes (**Fig. 3D**) compared to the dynamic secretomes (**Fig. 3E**). We performed Kyoto Encyclopedia of Genes and Genomes (KEGG) overrepresentation analysis of all static secretome DEPs using a dot plot and implicated several inflammatory pathways, including tumor necrosis factor (TNF), toll-like receptor (TLR), nuclear factor-kappa B (NF-κB), T and B cell receptor, interleukin, cytokine, and chemokine signaling (**Fig. S2A**). Supporting our YAP data in the present study, our IPA upstream regulator analysis found that Rho-associated protein kinase 1 (ROCK1) and other putative drivers of YAP activation (integrin beta-1; ITGB1, focal adhesion kinase; FAK, Ras-related C3 botulinum toxin substrate 1; Rac1) were predicted to be upregulated in the MRL/MpJ static secretome.

**Fig. 3.**
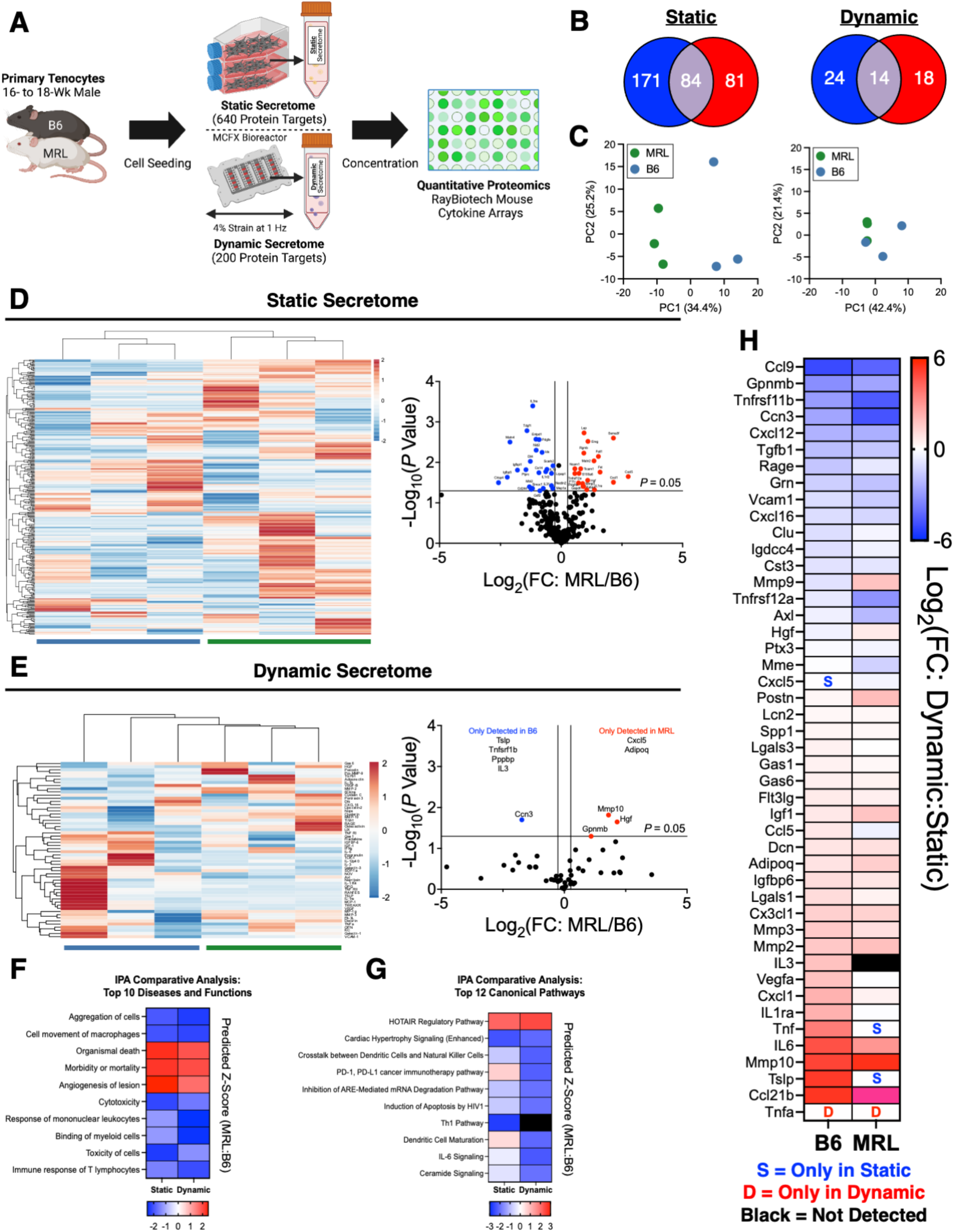
Quantitative proteomics reveal biological signatures enriched in the MRL/MpJ tenocyte secretome that are conserved across static and dynamic culture compared to those of B6. (**A**) Schematic illustration of primary mouse tenocyte secretome collection under static and dynamic culture conditions for RayBiotech quantitative proteomics analyses. (**B**) Breakdown of differentially expressed proteins (DEPs) upregulated in the B6 (in blue) and MRL/MpJ (in red) tenocyte secretomes. (**C**) Principal components analysis (PCA) showed distinct clustering of static secretome samples by genotype (left) but overlap of dynamic secretome samples (right). (**D and E**) Heat maps of Z-score hierarchical clustering based on Euclidean distance for rows and volcano plots of the static (D) and dynamic (E) secretomes. Blue and green bars underneath heat maps denote B6 and MRL/MpJ biological replicates, respectively. (**F to H**) Comparative ingenuity pathway analysis (IPA) of the static and dynamic secretomes predicted similar activation Z-score patterns for the top 10 diseases and functions (F) and canonical pathways (G) but contrasting activation states for 8 of 10 upstream regulators (H). (I) Heat map visualizing relative changes in protein concentration within each genotype for cytokine targets shared between the static and dynamic secretomes. Blue and red colors indicate annotations predicted to be activated or enriched in B6 or MRL/MpJ secretomes, respectively. Black boxes indicate annotations or cytokines that were not detected.

On the basis that tenocytes experience cyclic stretching due to dynamic tensile loading, we next conducted comparative analyses of the static and dynamic secretomes using IPA to determine canonical pathways, upstream regulators, and diseases and functions that are differentially regulated with mechanical stimulation. Surprisingly, mechanical stimulation had a minimal effect on the majority of diseases and functions (**Fig. 3F and Table S2**) and canonical pathways (**Fig. 3G**)—most of which are involved in inflammation and cell death—with similar Z-score activation patterns between the static and dynamic secretomes. In contrast, minimal agreement was found for upstream regulators with 8 of the top 10 annotations having contrasting activation states (**Fig. S1B**), suggesting molecular changes in B6 and MRL/MpJ tenocyte activity with mechanical stimulation that nonetheless lead to comparable cellular functions and outcomes. To assess the effects of mechanical stimulation on the secretion of soluble factors detected in both secretome datasets, we next compared the relative change in protein concentrations for 46 cytokines shared between the static and dynamic secretomes. Interestingly, we found that B6 and MRL/MpJ dynamic secretomes showed similar fold change expression for 37 cytokines (80.4% of total) compared to their respective static secretomes (**Fig. 3H**). Overall, these distinct paracrine signaling signatures between B6 and MRL/MpJ tenocytes regardless of loading modality illustrate further cell-intrinsic differences that distinguish scar-mediated and scarless tendon healing outcomes.

### Adult quiescent MRL/MpJ tenocytes undergo greater proliferative and apoptotic activity compared to B6 tenocytes

Altered cell cycle control (e.g., elevated G2/M accumulation) has been proposed as a potential characteristic of the regenerative MRL/MpJ cell phenotype (7), leading us to next characterize the growth kinetics and endogenous β-Galactosidase (β-Gal) activity, a marker for cellular senescence (i.e., irreversible cell cycle arrest), between B6 and MRL/MpJ tenocytes (**Fig. 4A and B**). B6 tenocytes showed a significantly higher percentage of β-Gal-positive/senescent cells at passage four (45.4%, *P* = 0.022) and five (60.7%, *P* < 0.0001) compared to passage three (25.3%; **Fig 4C**). In contrast, MRL/MpJ tenocytes displayed minimal cellular senescence with no significant changes in β-Gal expression with increasing passage numbers (0.3%, 1%, and 6%; **Fig. 4C**). These results were accompanied by a progressive decline in B6 tenocyte population doubling time (PDT) with serially passaging, whereas MRL/MpJ tenocyte PDT remained constant and was significantly lower than that of B6 cells at passage four (1.99-fold lower, *P* = 0.032) (**Fig. 4D**). Although growth rates were comparable at earlier passages, cumulative population doubling (CPD) at passage four was significantly greater in MRL/MpJ tenocytes compared to B6 tenocytes (1.32-fold higher, *P* = 0.0048). At each passage, both the percentage (*P* = 0.003 at P3, < 0.0001 at P4, < 0.0001 at P5) and number of senescent cells (*P* = 0.018 at P3, < 0.0001 at P4, < 0.0001 and P5; **Fig. S3A**) were significantly higher in B6 tenocytes compared to MRL tenocytes. The total cell number was not different between B6 and MRL/MpJ tenocytes (**Fig. S3B**). Collectively, these data demonstrate that MRL/MpJ tenocytes are more resistant to replicative senescence and exhibit superior expansion potential compared to B6 tenocytes. Given that growth arrest and resistance to pro-apoptotic stimuli are hallmarks of cellular senescence (30,31), we next performed immunohistochemical (IHC) analysis of Ki-67 and cleaved caspase-3, markers of cell proliferation and apoptosis respectively, of injured B6 and MRL/MpJ patellar tendons during the proliferative and remodeling phases of healing (**Fig. 4E**). As expected, MRL/MpJ tendons had significantly increased cell proliferation (**Fig. 4F**) and apoptosis (**Fig. 4G**) compared to B6 tendons at 1 wpi (Ki-67: *P* < 0.0001, caspase-3: *P* = 0.0032) and 4 wpi (Ki-67: *P* = 0.0027, caspase-3: *P* = 0.029), providing evidence that corroborates the relevance of our *in vitro* findings as potentially conserved mechanisms at play during the more complex *in vivo* healing environment.

**Fig. 4.**
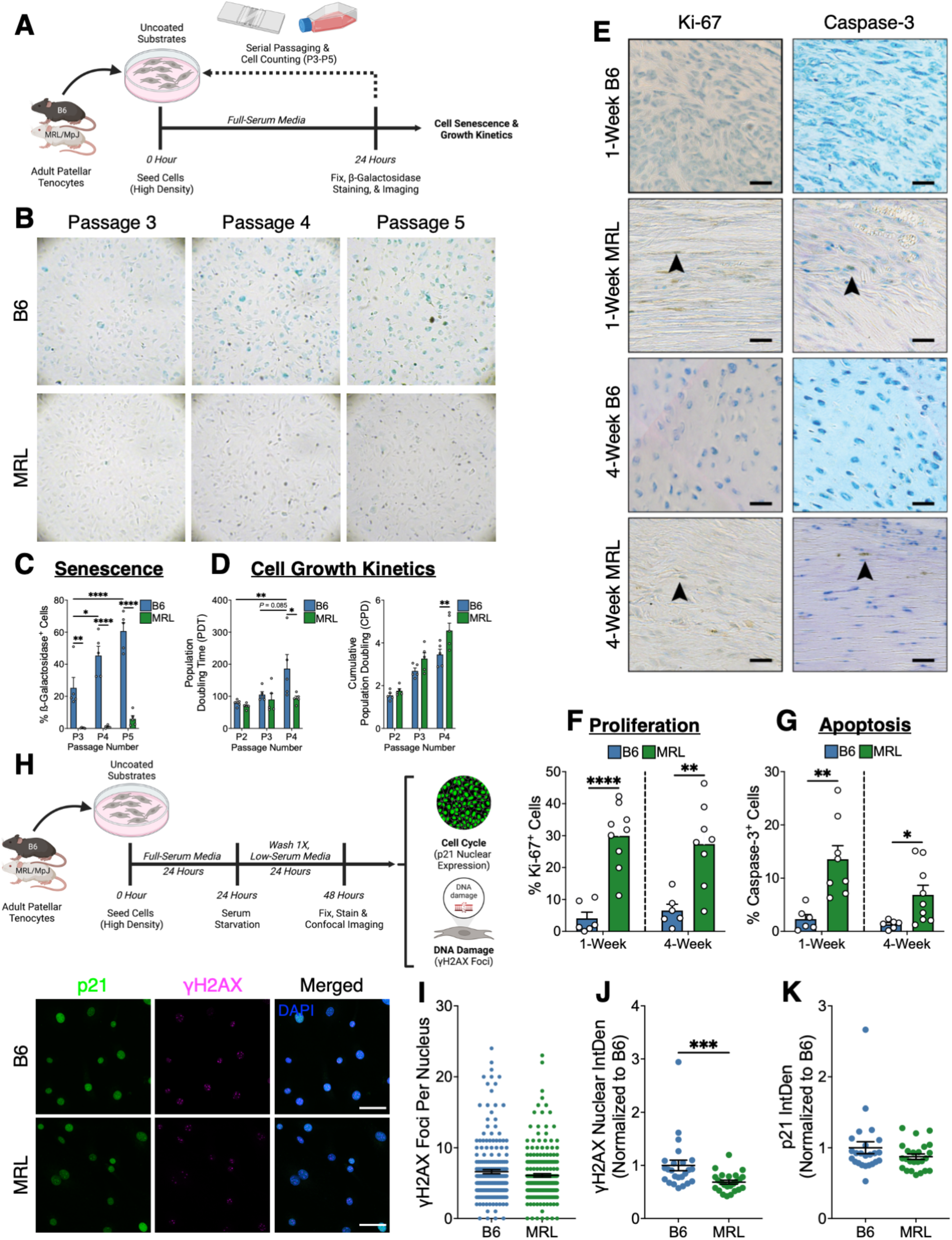
An adult population MRL/MpJ tenocytes undergo increased proliferative and apoptotic activity compared to B6 tenocytes. (**A**) Experimental design to characterize cellular senescence and the growth kinetics of B6 and MRL/MpJ tenocytes. (**B and C**) B6 tenocytes progressively accumulated ß-galactosidase (ß-gal) activity associated with cellular senescence in response to replicative stress induced by serial passages (P3-P5; B), whereas MRL/MpJ tenocytes maintained a non-senescent phenotype (C). (**D)** Population growth kinetics are diminished in B6 as compared to MRL/MpJ tenocytes at P5 as evidenced by significantly increased population doubling time (PDT) and decreased cumulative population doubling (CPD). (**E to G**) Immunohistochemical (IHC) staining of Ki-67 and cleaved caspase-3 found significantly increased cell proliferation (F) and apoptosis (G), respectively, in MRL/MpJ tendons compared to B6 tendons at 1 and 4 weeks (F) after acute injury *in vivo* (scale bar, 50 μm; arrowheads indicate positively-stained cells; all sections were counterstained with toluidine blue). (**H**) Experimental design to measure DNA damage and p21 expression in B6 and MRL/MpJ tenocytes. Immunofluorescence (IF) staining revealed similar intrinsic expression of p21(green) and DNA damage (indicated by γH2AX-positive foci, magenta) between B6 and MRL/MpJ tenocytes (scale bar, 100 µm). *\*P* < 0.05, \*\**P* < 0.01, \*\*\*\**P* < 0.0001. Significance was determined using a two-way analysis of variance (ANOVA) with post-hoc Bonferroni or Šidák for *in vitro* assays, or an unpaired Student’s t-test when comparing genotypes at each timepoint for IHC data. Representative images shown for all staining experiments.

The p53/p21 signaling axis is thought to play a critical role in MRL/MpJ tissue regeneration wherein the absence of p21 promotes senescence, hyperproliferation, and the DNA damage response (32). Notably, p21^-/-^ knockout and MRL/MpJ mice are capable of regenerating full-thickness ear and articular cartilage defects (32,33). To determine whether reduced p21 expression and endogenous DNA damage are associated with MRL/MpJ tenocyte behavior *in vitro*, we stained B6 and MRL/MpJ cells for endogenous p21 and γH2AX. Contrary to reports of linking appendage regeneration to a lack of p21 expression in MRL/MpJ ear fibroblasts(32), we detected endogenous p21 activity in MRL/MpJ tenocytes (**Fig. 4H**). There were no differences in the number of γH2AX-positive foci that mark DNA double-stranded breaks (**Fig. S4A**) or p21 expression (**Fig. S4B**), indicating that MRL/MpJ tenocyte quiescence and proliferation may be regulated independently of p21 and DNA damage repair mechanisms.

After elucidating key biological processes underlying MRL/MpJ tenocyte behavior, we interrogated the biochemical milieu of the local MRL/MpJ tendon environment by culturing B6 and MRL/MpJ tenocytes using a combination of their dPECM-coated substrates (12) and the secretome.

### B6 and MRL/MpJ tenocytes treated with MRL/MpJ-derived proteins attain comparable single-cell morphology and cell-ECM interactions

We have previously shown that dPECM harvested from MRL/MpJ tendon at 1 wpi enhance B6 tenocyte cell morphology and proliferation *in vitro*. Additionally, comparative analyses results in IPA showed similar biological annotations between the static and dynamic secretomes, motivating the use of the static secretome in subsequent experiments for practicality. Accordingly, we investigated three experimental conditions: (*i*) a scar-mediated healing group of B6 tenocytes treated with B6-derived components (i.e., dPECM and secretome) (referred to as ‘BB’), (*ii*) a therapeutic group of B6 tenocytes treated with MRL/MpJ-derived components (referred to as ‘BM’), and (*iii*) a positive control group of MRL/MpJ tenocytes treated with MRL/MpJ-derived components (referred to as ‘MM’).

Strikingly, B6 and MRL/MpJ tenocytes at a low cell density treated with MRL/MpJ-derived components displayed identical single-cell morphology, FA formation, and cellular protrusion patterns (**Fig. 5A and B**). In contrast, BB exhibited significantly higher cell circularity and lower perimeter, number of FAs and total FA area, and protrusion parameters (percentage of cells with protrusions, number of protrusions, total protrusion length, total protrusion length normalized to cell perimeter) compared to BM and MM (**Fig. 5D-F**). Additionally, B6 tenocytes treated with MRL/MpJ-derived components exhibited significantly higher αSMA expression compared to B6 tenocytes treated with B6-derived components (*P* = 0.016) or at baseline (*P* < 0.0001) (**Fig. 5G and H**), suggesting that myofibroblast activity is transiently increased but does not remain elevated with extended culture periods. Interestingly, B6-derived components elicited little or no effect with BB being largely comparable to untreated B6 tenocytes. Supporting our hypothesis, these findings demonstrate that MRL/MpJ-derived components reprogram cell morphology and amplify mechanotransduction pathways towards MRL/MpJ tenocyte behavior.

**Fig. 5.**
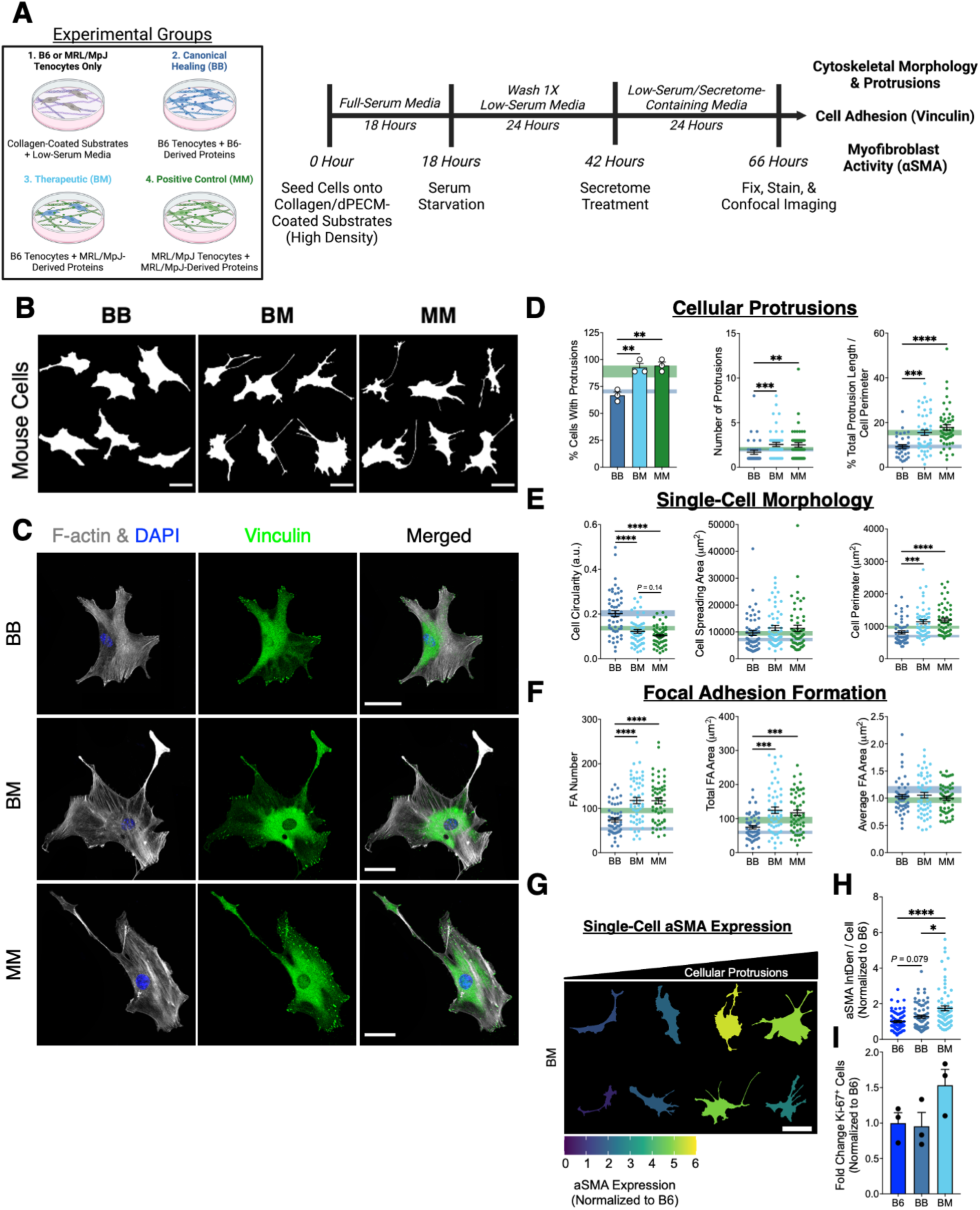
B6 and MRL/MpJ tenocytes treated with a combination of MRL/MpJ-derived decellularized provisional ECM and secretome achieve comparable morphologies and cellular mechanotransduction. (**A**) Experimental design and *in vitro* culture model of single-cell experiments for tenocytes cultured on dPECM with tenocyte-sourced secretome (henceforth referred to as ‘components’). (**B**) Representative cell outlines of B6 and MRL/MpJ tenocytes cultured with B6 or MRL/MpJ-derived components (scale bar, 100 μm) to visualize cytoskeletal morphology and cellular protrusions. (**C**) IF staining of F-actin (grey) and vinculin (green) of treated B6 and MRL/MpJ tenocytes. (**D to F**) Cellular protrusion characteristics (D), single-cell morphological parameters (E), and focal adhesion profiles (F) are identical between B6 and MRL/MpJ tenocytes treated with MRL/MpJ-derived components and significantly different from B6 tenocytes treated with B6-derived components. (**g**) Representative cell outlines of B6 tenocytes cultured with MRL/MpJ-derived components (scale bar, 200 μm). Cells with greater protrusion numbers and total protrusion lengths were associated with increased myofibroblast activity as visualized by dark blue colors indicating higher αSMA expression (relative to B6 mean). (**H and I**) MRL/MpJ-derived components promoted significantly greater αSMA expression (H) in B6 tenocytes but not cell proliferation (I) compared to B6-derived components or B6 cells cultured on collagen I-coated substrates. \**P* < 0.05, \*\**P* < 0.01, \*\*\**P* < 0.001, \*\*\*\**P* < 0.0001. Significance was determined using a one-way Welch’s ANOVA with post-hoc Games-Howell for cell morphology and focal adhesion data. For cellular protrusion data, significance was determined using a one-way ANOVA with post-hoc Tukey (percentage of cells with protrusions) or Kruskal-Wallis test with post-hoc Dunn’s (number of protrusions and normalized total protrusion length). Representative images shown for all staining experiments.

We next assessed the capacity of MRL/MpJ-derived components to enhance cytoskeletal organization, ECM deposition, and Cx-43 formation in confluent B6 tenocytes (**Fig. 6A**). To minimize the influence of serum proteins and confounding effects from differences in total cell number due to increased MRL/MpJ tenocyte proliferation, cells were seeded at a high density and subsequently maintained in low-serum media with or without exogenous secretome supplementation. Moreover, to definitively ascertain whether there is a combined benefit of administering the MRL/MpJ-derived components, we also cultured B6 tenocytes with MRL/MpJ dPECM or secretome alone with cells cultured in only low-serum media served as baseline controls. Consistent with our previous baseline B6 and MRL/MpJ comparisons under culture in full-serum media, MRL/MpJ tenocytes showed greater baseline F-actin alignment (*P* = 0.0003), fibronectin deposition (1.20-fold higher, *P* = 0.095), collagen I deposition (1.15-fold higher, *P* = 0.0029), and no difference in αSMA expression compared to B6 tenocytes (**Fig. 6B**). In contrast with our previous data in full-serum media, MRL/MpJ tenocytes exhibited significantly lower baseline Cx-43 expression than B6 tenocytes under low-serum conditions (1.96-fold lower, *P* < 0.0001). Supporting our hypothesis, B6 and MRL/MpJ tenocytes treated with MRL/MpJ-derived components showed comparable F-actin alignment and relative changes in fibronectin deposition that were significantly higher than that of B6 tenocytes treated with B6-derived components (**Fig. 6C and D**). BM also showed greater collagen I deposition compared to BB (*P* = 0.10) (**Fig. S6A**). Due to restricted ECM deposition under low-serum conditions, fibronectin alignment could not be quantified. Contrary to our hypothesis, BB and BM had comparable Cx-43 expression that were significantly lower than MM. However, these discrepancies are largely attributed to a nearly two-fold reduction in Cx-43 expression in MRL/MpJ at baseline, as evidenced by comparable staining profiles between cells that received any combination of mouse-derived components. This data suggest that these tenocytes upregulate Cx-43 expression in response to mitogenic stimulation. There were no differences in αSMA expression between all treatment groups. Further supporting our hypothesis, F-actin alignment and Cx-43 levels were significantly higher in B6 tenocytes treated with MRL/MpJ-derived components compared to dPECM (*P* = 0.0031 for F-actin and *P* < 0.0001 for Cx-43) or secretome (*P* = 0.031 for F-actin and *P* = 0.002 for Cx-43) alone (**Fig. 6E**), confirming the combined benefit of administering the MRL/MpJ-derived components.

**Fig. 6.**
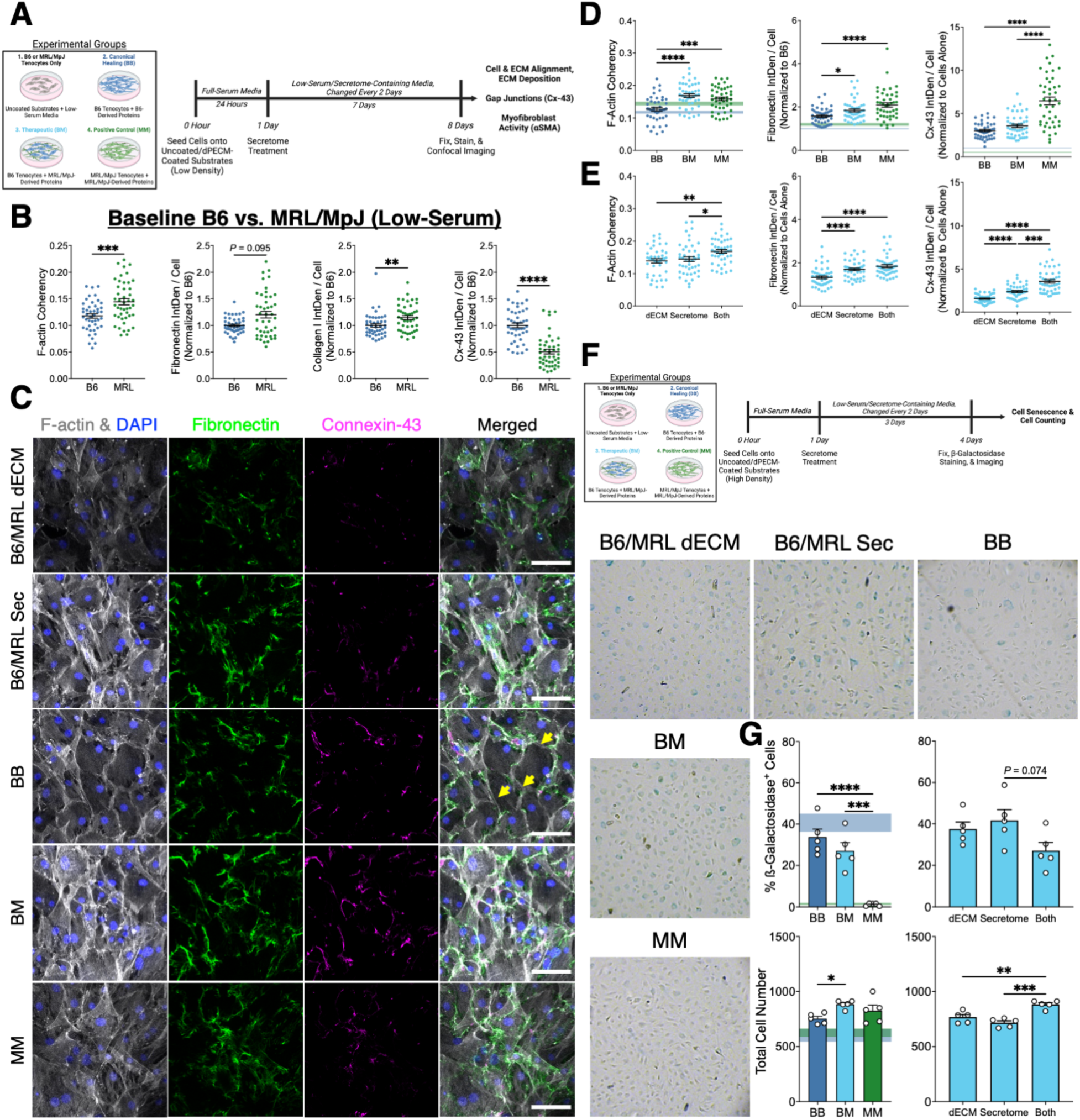
Combination of MRL/MpJ-derived components modulate confluent B6 tenocytes toward an MRL/MpJ-like regenerative phenotype *in vitro*. (**A**) Experimental design and *in vitro* culture model of confluent cell experiments for tenocytes cultured with mouse-derived components for 7 days. (**B**) MRL/MpJ tenocytes maintained significantly greater F-actin alignment and ECM deposition than B6 tenocytes under serum-deprived culture conditions. However, Cx-43 levels were significantly downregulated in MRL/MpJ tenocytes compared to B6 tenocytes. (**C**) IF staining of F-actin (grey), fibronectin (green), and Cx-43 (magenta) of confluent B6 and MRL/MpJ tenocytes treated with only MRL/MpJ dPECM, only MRL/MpJ secretome, or a combination of mouse-derived components for 7 days (scale bar, 100 μm). A polygonal, epithelial cell-like morphology was observed in B6 tenocytes treated with B6-derived components (indicated by yellow arrowheads). (**D**) B6 and MRL/MpJ tenocytes treated with MRL/MpJ-derived components exhibited comparable F-actin alignment and fibronectin deposition that were significantly greater than those of B6 tenocytes treated with B6-derived components. However, both treated B6 tenocyte groups had significantly lower Cx-43 levels compared to MRL/MpJ tenocytes treated with MRL/MpJ-derived components. (**E**) There was a combined benefit of treating B6 tenocytes with both the MRL/MpJ dPECM and secretome on F-actin alignment, fibronectin deposition, and Cx-43 levels compared to either component alone. (**F**) Experimental design to quantify cellular senescence in tenocytes cultured with mouse-derived components for 3 days. MRL/MpJ components were unable to attenuate the onset of B6 tenocyte senescence/accumulation of ß-gal activity after 48 hours of treatment. (**G**) B6 tenocytes treated with MRL/MpJ-derived components displayed a smaller proportion of senescent cells due to significantly increased total cell number compared to either component alone as well as B6 tenocytes treated with B6-derived components. \**P* < 0.05, \*\**P* < 0.01, \*\*\**P* < 0.001, \*\*\*\**P* < 0.0001. Significance was determined using a one-way ANOVA with post-hoc Tukey. Representative images shown for all staining experiments.

Lastly, we determined whether MRL/MpJ-derived components could promote cell proliferation and attenuate the onset of cellular senescence by treating B6 tenocytes cultured at a moderate density for 3 days and staining for ß-gal activity (**Fig. 6F**). There was no difference in cell number between B6 and MRL/MpJ tenocytes at baseline or between cells treated with MRL/MpJ-derived components. Supporting our hypothesis, compared with BB, BM showed significantly increased total cell number (1.17-fold higher, *P* = 0.045). In addition, total cell number was significantly higher in B6 tenocytes treated with MRL/MpJ-derived components compared to dPECM (*P* = 0.036) or secretome (*P* = 0.0002) alone (**Fig. 6G**). These results are consistent with previous observations that an early increase in tenocyte proliferation is a hallmark of MRL/MpJ tendon healing(10,12). However, contrary to our hypothesis, there was no difference in the number or percentage of senescent cells between BB and BM. B6 tenocytes treated with only secretome exhibited less senescence overall compared to BM (1.15-fold lower, *P* = 0.074), but no differences were found in the number of senescent cells between all B6 tenocyte groups. Altogether, these findings demonstrate that MRL/MpJ-derived components potently augment B6 tenocytes towards pro-regenerative MRL/MpJ-like behavior despite the persistence of a senescent subpopulation of cells.

Collectively, our data provide compelling evidence that the combined administration of MRL/MpJ-derived components modulates B6 tenocytes toward an MRL/MpJ-like cell phenotype. However, there are important considerations in harnessing these biological cues for therapeutic delivery. First, the tendon ECM and secretome each contain thousands of biomolecules (34,35), which can preclude their feasibility to scale up for preclinical testing in animal models. Second, it remains unclear whether these hallmarks of MRL/MpJ tenocyte behavior and the therapeutic utility of MRL/MpJ-derived components are conserved across other mammalian species. To address these points, we sought to develop a panel of recombinant proteins using our quantitative proteomics datasets. The final objective of this study was to assess the functional activity of B6-derived components, MRL/MpJ-derived components, and the recombinant protein therapeutic *in vitro* on patellar tenocytes derived from scar-mediated healing Sprague-Dawley (SD) rats. We hypothesized that the recombinant protein therapeutic will recapitulate the effects of the MRL/MpJ-derived components on rat tenocyte behavior *in vitro* and that these changes will be consistent with our mouse data.

### Recombinant protein therapeutic promotes regenerative MRL/MpJ-like tenocyte behavior across species

Mouse and rat models closely recapitulate human tendon anatomy and physiology (36,37), supporting the use of rodent tenocytes to investigate the proof-of-concept translational capacity of our recombinant protein therapeutic. However, despite the prevalent use of rodent models for tendon research, a comparative evaluation of baseline mouse and rat tenocytes behavior is lacking. Accordingly, we first cultured B6 and SD patellar tenocytes on collagen I-coated substrates and analyzed their single-cell morphology. There were no differences in any morphological (cell circularity, spreading area, perimeter), cellular protrusion parameters (percentage of cells with protrusions, number of protrusions, total protrusion length), or αSMA expression between B6 and SD tenocytes (**Fig. S7A and B**)

A candidate panel of 29 recombinant proteins (10 structural and 19 soluble, 12 of which were from the secretome) was identified from a systemic screen of our quantitative proteomics data (**Table 1**). However, a preliminary set of 17 recombinant proteins (indicated in bold) were employed in the present study for our protein therapeutic (**Fig. 7A**) due to several proteins being on backorder or lacking commercial availability at the time of investigation. To mimic our combined dPECM and secretome culture system with the recombinant protein therapeutic, structural proteins were sequentially adhered onto a collagen I-coated substrate starting with acid-solubilized proteins followed by those reconstituted in sterile 1X PBS. Soluble proteins were supplemented into low-serum media. All protein concentrations used in this study were determined from published reference values for *in vitro* administration. We then investigated three experimental conditions: (*i*) a scar-mediated healing group of SD tenocytes treated with B6-derived components (referred to as ‘RB’), (*ii*) a positive control group of SD tenocytes treated with MRL/MpJ-derived components (referred to as ‘RM’), and (*iii*) the therapeutic group of SD tenocytes treated with the recombinant protein therapeutic (referred to as ‘RP’). For the B6-derived components, B6 uninjured dECM was used as the substrate coating instead of the dPECM to match the conditions analyzed via mass spectrometry in our published *in vivo* study (16).

**Fig. 7.**
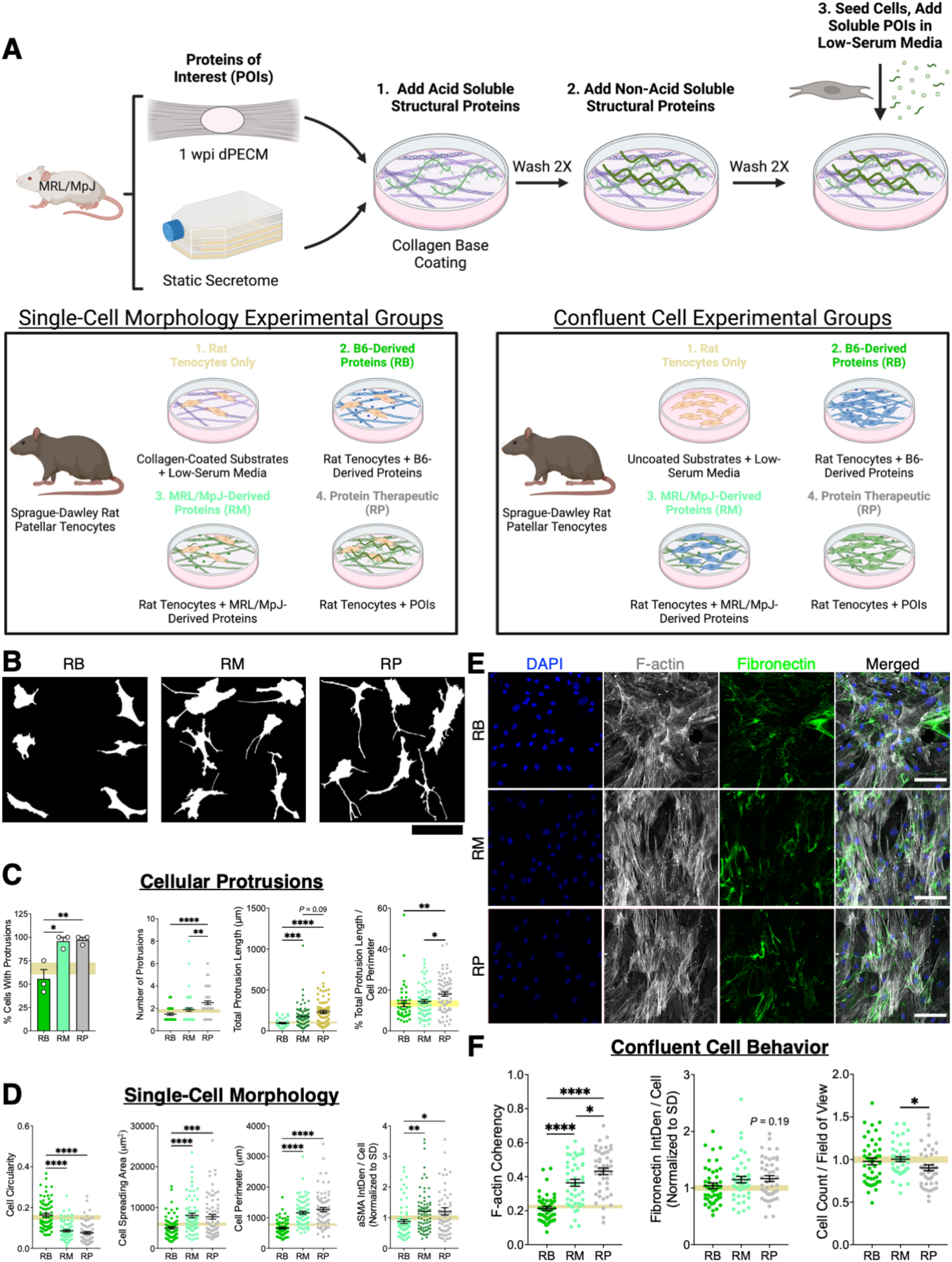
A recombinant protein therapeutic comprised of MRL/MpJ-enriched structural and soluble factors phenocopies MRL/MpJ cell behavior in Sprague-Dawley rat tenocytes *in vitro*. (**A**) Experimental design of recombinant protein therapeutic development and *in vitro* culture model for single-cell and confluent cell experiments. (**B**) Representative cell outlines of Sprague-Dawley (SD) rat tenocytes cultured with B6 or MRL/MpJ-derived components or the recombinant protein therapeutic (scale bar, 100 μm) to visualize cytoskeletal morphology and cellular protrusions. (**C and D**) SD tenocytes treated with MRL/MpJ-derived components and the recombinant protein therapeutic exhibited identical (or superior in the protein therapeutic) cellular protrusion characteristics (C) and single-cell morphology (D) that were significantly different from SD tenocytes treated with B6-derived components. (**E**) IF staining of F-actin (grey) and fibronectin (green) of confluent SD tenocytes treated with mouse-derived components or the recombinant protein therapeutic (scale bar, 100 μm). (**F**) SD tenocytes treated with the recombinant protein therapeutic showed significantly higher F-actin alignment and similar fibronectin deposition compared to SD tenocytes treated with mouse-derived components. Total cell number was significantly reduced by the recombinant protein therapeutic relative to the MRL/MpJ-derived components. \**P* < 0.05, \*\**P* < 0.01, \*\*\**P* < 0.001, \*\*\*\**P* < 0.0001. Significance was determined using a one-way Welch’s ANOVA with post-hoc Games-Howell for cell morphology, total protrusion length, and total cell number data. For cellular protrusion and fibronectin deposition data, significance was determined using a one-way ANOVA with post-hoc Tukey (percentage of cells with protrusions) or Kruskal-Wallis test with post-hoc Dunn’s (number of protrusions, normalized total protrusion length, normalized fibronectin integrated density). Representative images shown for all staining experiments.

**Table 1.**
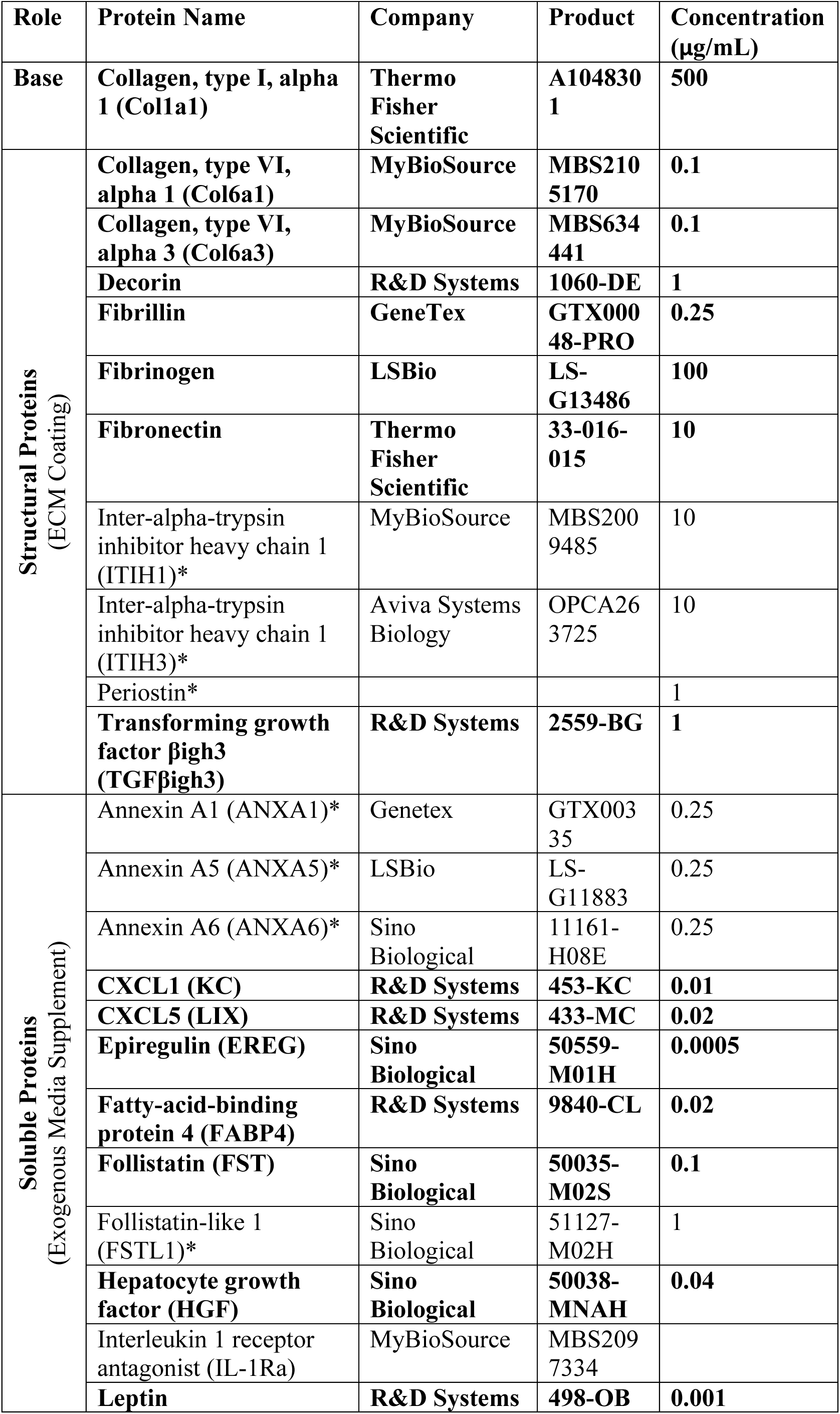
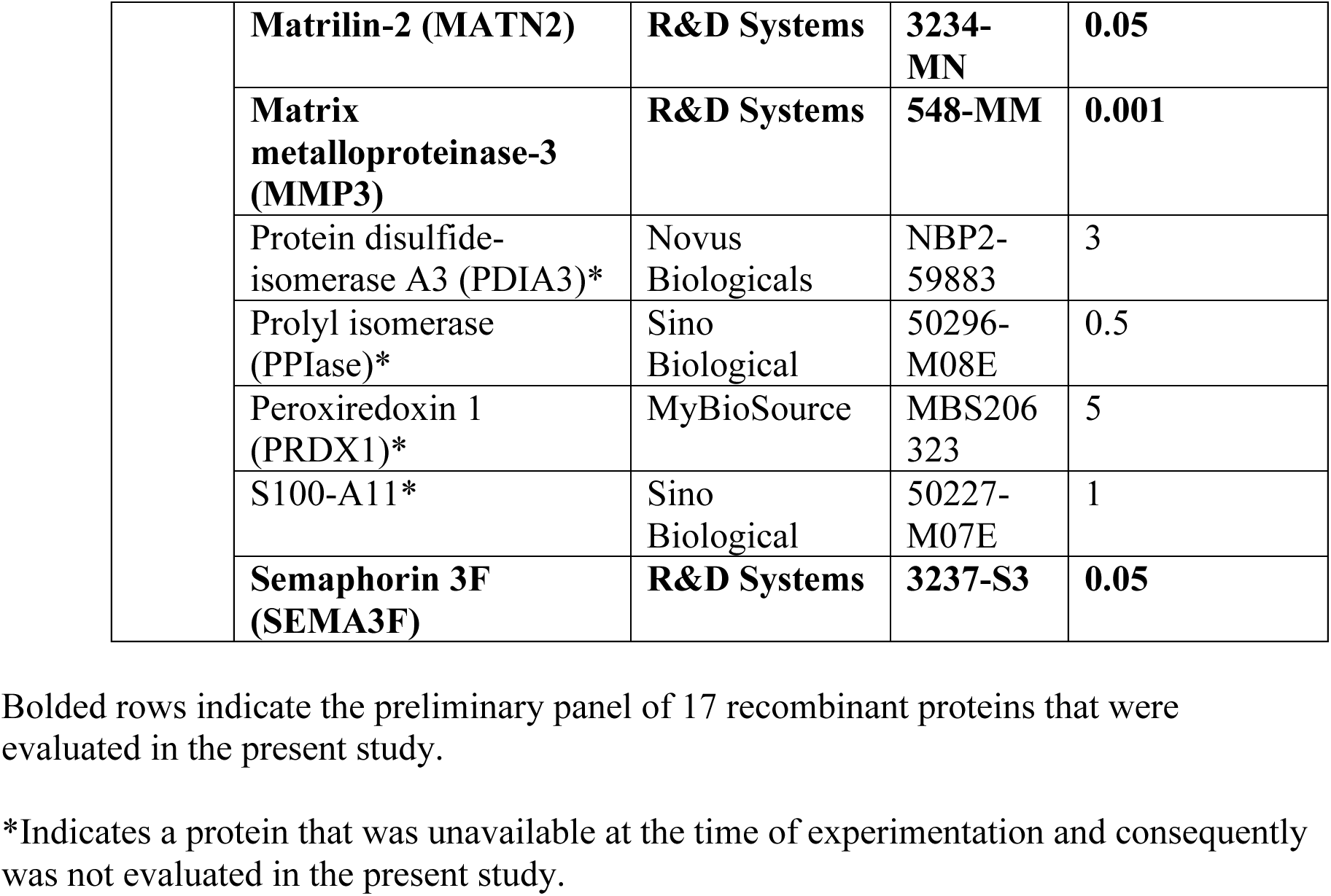
Structural and soluble recombinant proteins enriched in the MRL/MpJ provisional extracellular matrix and secretome identified using quantitative proteomics.

**Table 2.**
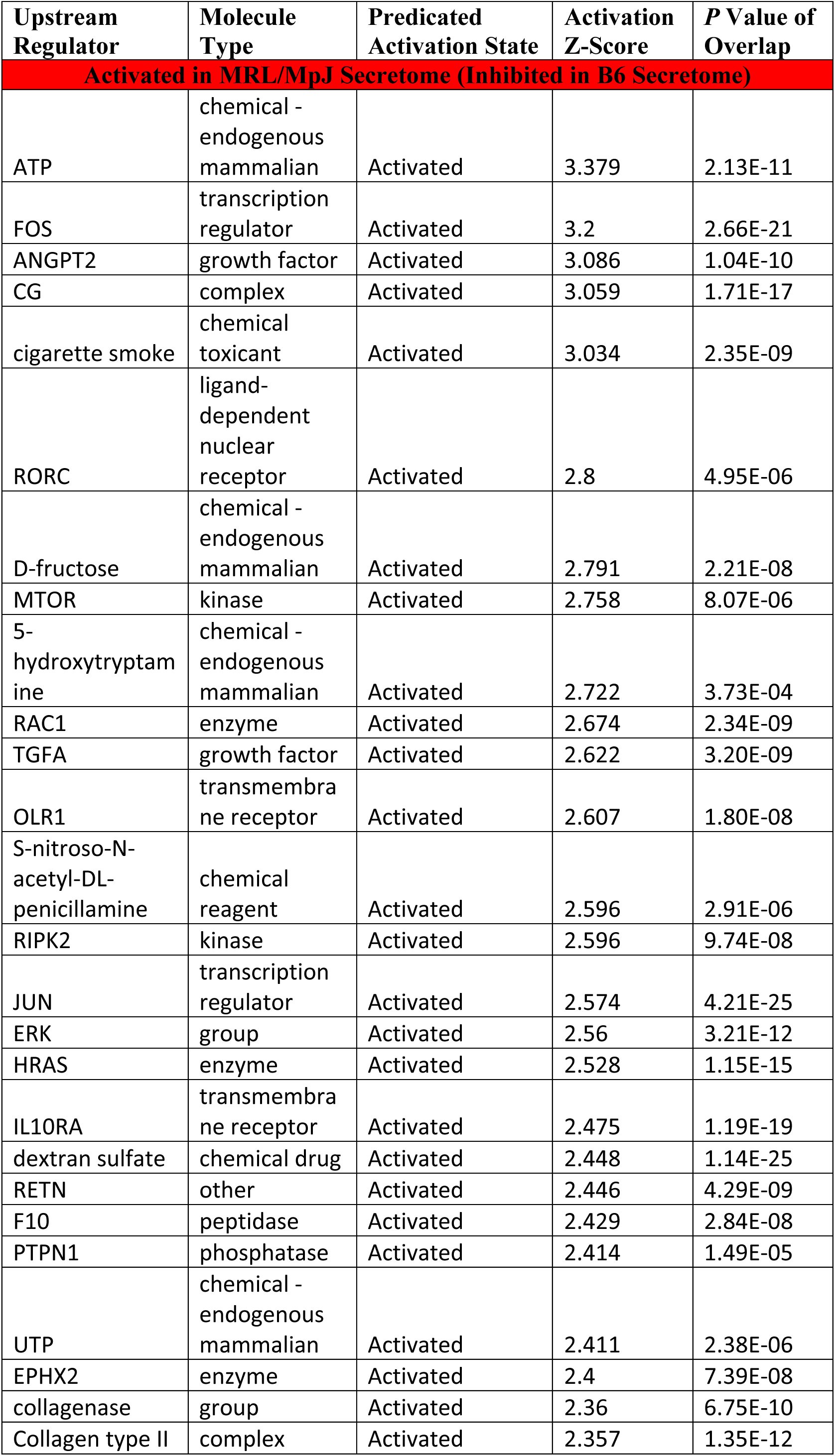
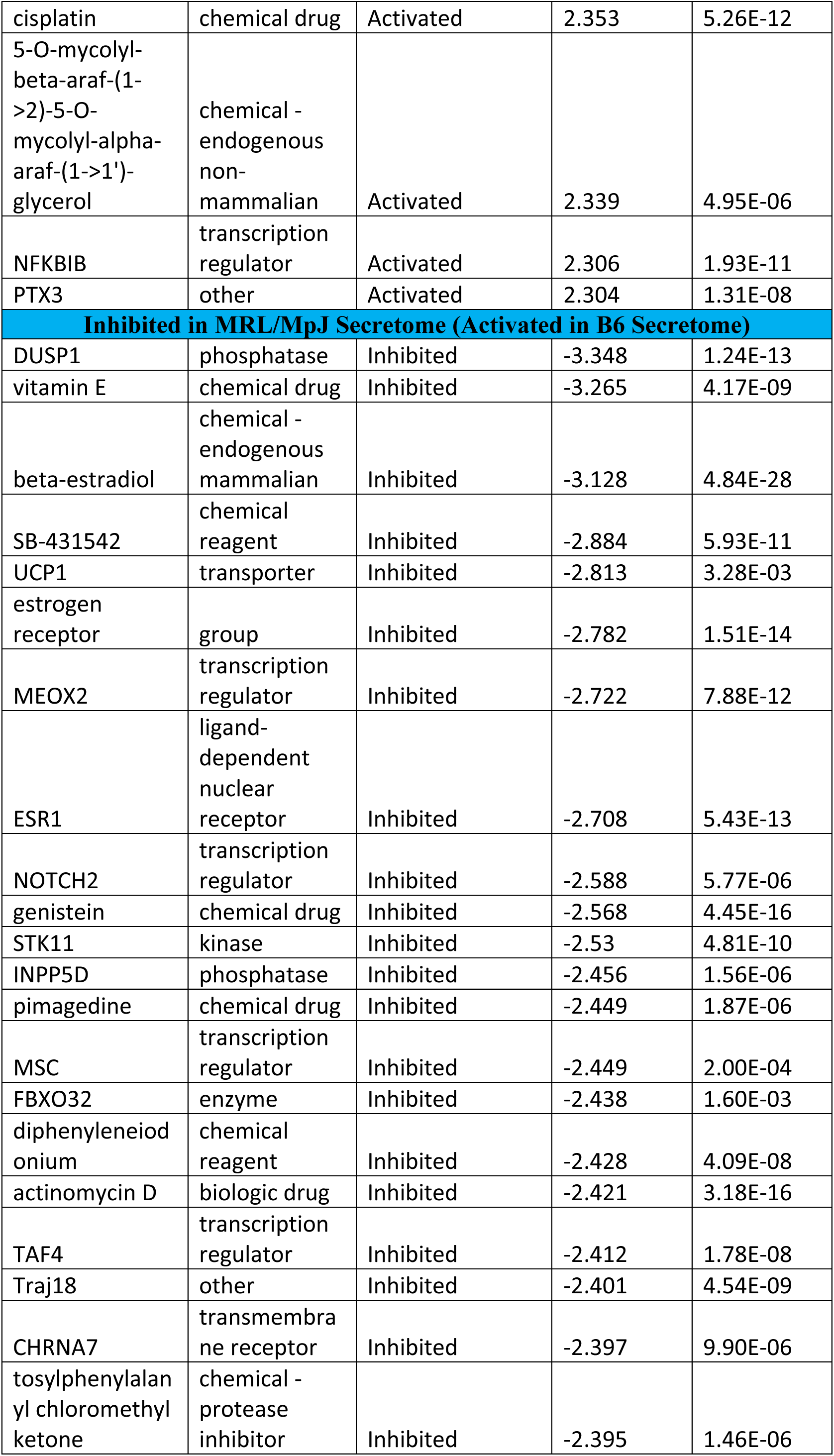
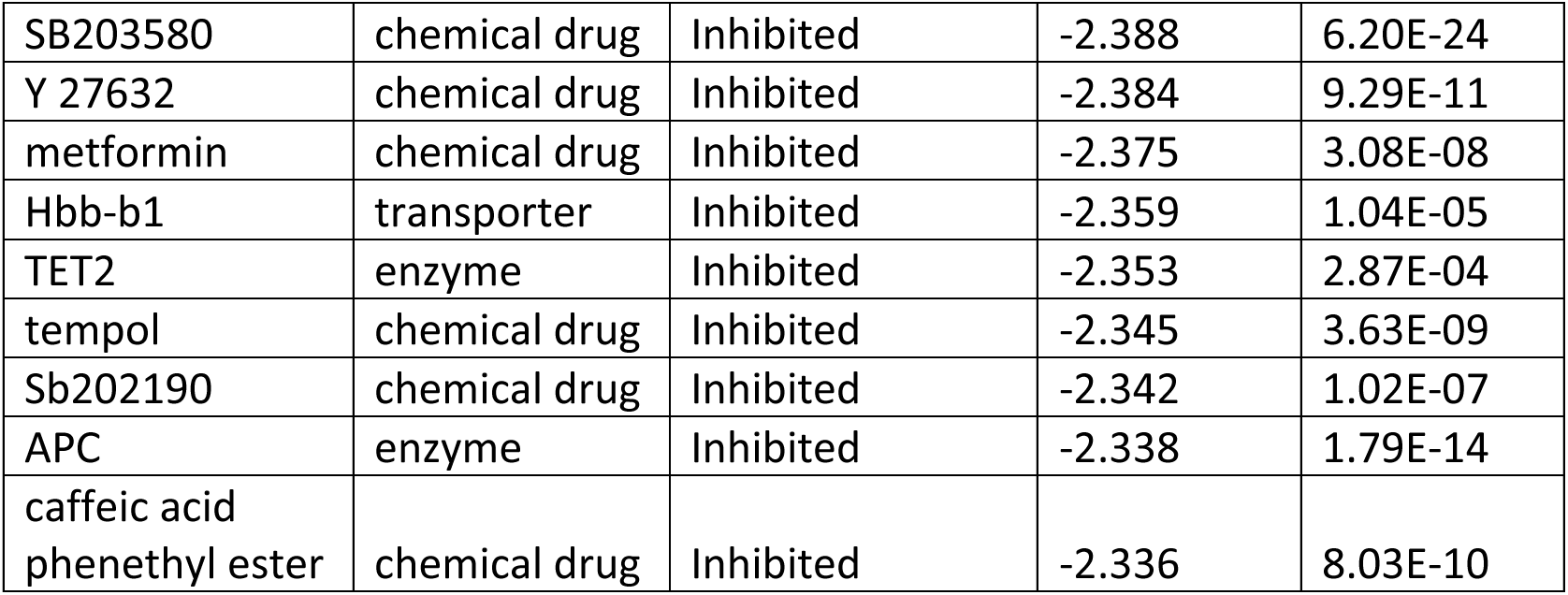
Top 30 activated and inhibited ingenuity pathway analysis upstream regulators from proteomic analyses of the static secretome.

**Table 3.**
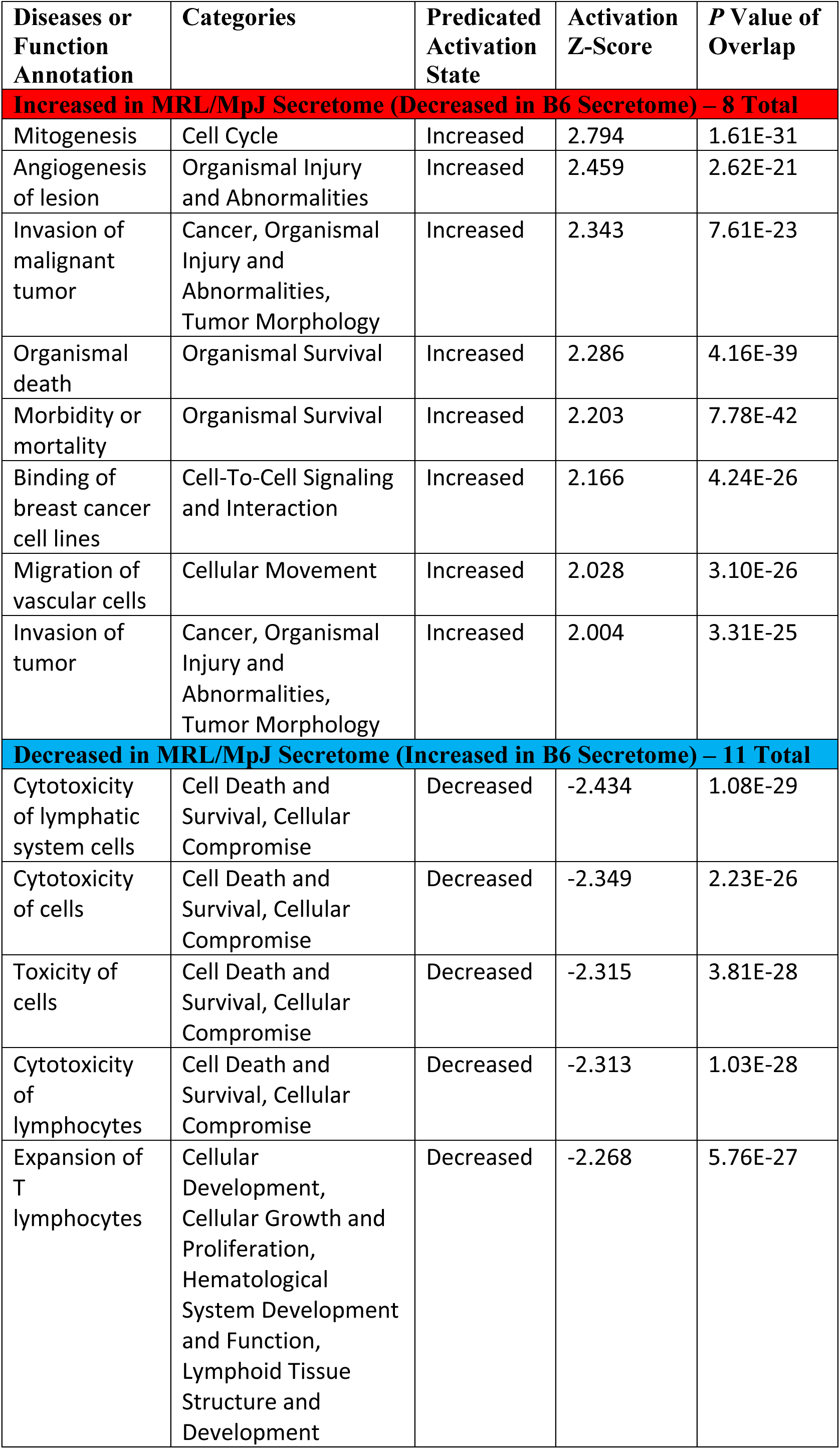
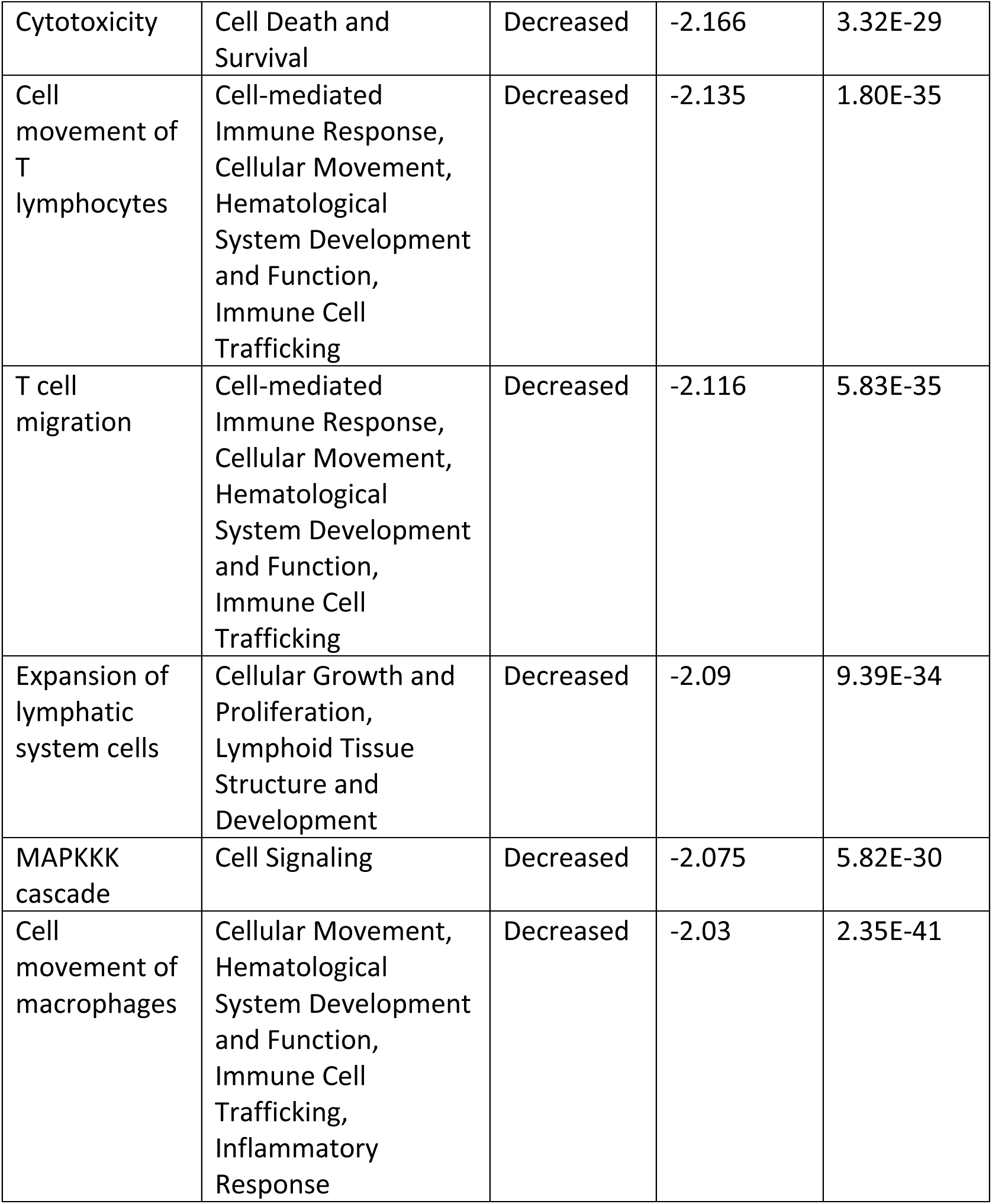
Increased and decreased ingenuity pathway diseases and functions from proteomic analyses of the static secretome.

| Diseases or Function Annotation | Categories | Predicated Activation State | Activation Z-Score | P Value of Overlap |
| --- | --- | --- | --- | --- |
| **Increased in MRL/MpJ Secretome (Decreased in B6 Secretome) – 8 Total** |  |  |  |  |
| Mitogenesis | Cell Cycle | Increased | 2.794 | 1.61E-31 |
| Angiogenesis of lesion | Organismal Injury and Abnormalities | Increased | 2.459 | 2.62E-21 |
| Invasion of malignant tumor | Cancer, Organismal Injury and Abnormalities, Tumor Morphology | Increased | 2.343 | 7.61E-23 |
| Organismal death | Organismal Survival | Increased | 2.286 | 4.16E-39 |
| Morbidity or mortality | Organismal Survival | Increased | 2.203 | 7.78E-42 |
| Binding of breast cancer cell lines | Cell-To-Cell Signaling and Interaction | Increased | 2.166 | 4.24E-26 |
| Migration of vascular cells | Cellular Movement | Increased | 2.028 | 3.10E-26 |
| Invasion of tumor | Cancer, Organismal Injury and Abnormalities, Tumor Morphology | Increased | 2.004 | 3.31E-25 |
| **Decreased in MRL/MpJ Secretome (Increased in B6 Secretome) – 11 Total** |  |  |  |  |
| Cytotoxicity of lymphatic system cells | Cell Death and Survival, Cellular Compromise | Decreased | -2.434 | 1.08E-29 |
| Cytotoxicity of cells | Cell Death and Survival, Cellular Compromise | Decreased | -2.349 | 2.23E-26 |
| Toxicity of cells | Cell Death and Survival, Cellular Compromise | Decreased | -2.315 | 3.81E-28 |
| Cytotoxicity of lymphocytes | Cell Death and Survival, Cellular Compromise | Decreased | -2.313 | 1.03E-28 |
| Expansion of T lymphocytes | Cellular Development, Cellular Growth and Proliferation, Hematological System Development and Function, Lymphoid Tissue Structure and Development | Decreased | -2.268 | 5.76E-27 |
| Cytotoxicity | Cell Death and Survival | Decreased | -2.166 | 3.32E-29 |
| Cell movement of T lymphocytes | Cell-mediated Immune Response, Cellular Movement, Hematological System Development and Function, Immune Cell Trafficking | Decreased | -2.135 | 1.80E-35 |
| T cell migration | Cell-mediated Immune Response, Cellular Movement, Hematological System Development and Function, Immune Cell Trafficking | Decreased | -2.116 | 5.83E-35 |
| Expansion of lymphatic system cells | Cellular Growth and Proliferation, Lymphoid Tissue Structure and Development | Decreased | -2.09 | 9.39E-34 |
| MAPKKK cascade | Cell Signaling | Decreased | -2.075 | 5.82E-30 |
| Cell movement of macrophages | Cellular Movement, Hematological System Development and Function, Immune Cell Trafficking, Inflammatory Response | Decreased | -2.03 | 2.35E-41 |

Based on our results that cell morphology is preserved across B6 and SD tenocytes, we were able to definitively ascertain the effects of our recombinant protein therapeutic on SD tenocyte behavior. Compared with MRL/MpJ-derived components, the recombinant protein therapeutic increased the number of protrusions (*P* = 0.0088) and total protrusion length (*P* = 0.09, 0.041 when normalized to cell perimeter) (**Fig. 7B and C**). Additionally, there were no differences in the number of protrusions or total protrusion coverage (normalized to cell perimeter) between RB, RM, or baseline. As expected, SD tenocytes cultured with the recombinant protein therapeutic and MRL/MpJ-derived components exhibited comparable cell circularity, spreading area, and αSMA expression (**Fig. 7D**). Supporting our hypotheses, RM and RP exhibited significantly lower circularity and higher perimeter, spreading area, αSMA expression, percentage of cells with protrusions, and total protrusion length compared to RB. Taken together, these results establish that the morphological features of MRL/MpJ tenocyte can be achieved in SD tenocytes through the selective delivery of key proteins. Further supporting our hypotheses, SD tenocytes treated with the recombinant protein therapeutic and MRL/MpJ-derived components showed comparable F-actin alignment and were significantly higher than that of SD tenocytes treated with B6-derived components (*P* < 0.0001 for both) (**Fig. 7E and F**). Moreover, RP displayed significantly higher F-actin alignment compared to RM (*P* = 0.02). Contrary to our hypothesis, while the recombinant protein therapeutic (*P* = 0.09) and MRL/MpJ components (*P* = 0.37) increased fibronectin deposition compared to the baseline control, these changes were not statistically significant. However, this may potentially be attributed to the recombinant protein therapeutic only containing 17 out of the 29 identified candidates. Lastly, we found significantly lower total cell number in RP compared to RM (9.8% lower; *P* = 0.04), suggesting that the recombinant protein therapeutic induces apoptosis that leads to more effective cytoskeletal alignment consistent with increased cell turnover during MRL/MpJ scarless tendon healing *in vivo*.

## Discussion

The widespread benefits of using tissue-specific dECM (38,39) and mesenchymal stem/progenitor cell-sourced secretome (35,40) to stimulate tissue repair and regeneration have been thoroughly established. However, there has been little exploration on the effects of these components together despite their capability to support complex cell-ECM and cell-cell interactions. Supporting the use of this combined biologic approach, our study integrated the structural constituents and cell-secreted soluble factors enriched in the MRL/MpJ tendon dPECM and secretome to modulate scar-mediated healing tenocytes towards an MRL/MpJ-like phenotype *in vitro*. These findings also provide mechanistic insight into the biological processes underlying regenerative MRL/MpJ tenocyte behavior and establish *in vitro* metrics grounded in physiologically relevant *in vivo* acute injury outcome measures. A major hurdle to harnessing the MRL/MpJ dPECM and secretome for therapeutic applications is that they each contain thousands of biomolecules, most of which may not be essential in delineating scar-mediated and regenerative healing, which presents challenges in translating these components for larger animal testing. Accordingly, our study isolated a panel of candidate protein regulators of mammalian tendon regeneration that exerts interspecies utility by enhancing scar-mediated healing rat tenocyte activity. Our data showed that rat tenocytes treated with the recombinant protein therapeutic outperformed the MRL/MpJ-derived components as evidenced by increased cellular protrusion formation and cytoskeletal alignment. These findings indicate that our approach of selectively administering recombinant proteins is functionally advantageous, which may be attributed to limiting unnecessary protein-protein interactions and instead achieving more targeted modulation of the molecular processes and downstream effectors that regulate MRL/MpJ cell behavior.

Constitutive stabilization of HIF-1α expression and p21 downregulation have both also been shown to control ear wound regeneration in adult MRL/MpJ mice (7,41). In contrast, multilineage differentiation, pluripotency markers, and HIF-1α expression were not observed in this study. We also found no difference in p21 expression between B6 and MRL/MpJ tenocytes consistent with findings reported by Lalley *et al* (10). Therefore, our data implicate resident tenocyte or progenitor cell populations as key mediators of MRL/MpJ tendon healing outcomes. Supporting our proposed paradigm, Grinstein *et al*. (42) recently identified a latent population of *Axin2^+^*tenocytes in the midsubstance of mouse and human tendons that is recruited to the injury site, proliferate, and transiently adopt injury responsive states to centrally organize the healing response. Our results illustrated that rodent tenocytes treated with MRL/MpJ-derived components or our recombinant protein therapeutic are robustly reprogrammed *in vitro* to attain morphological characteristics identical to those of MRL/MpJ tenocytes. This observation generates an exciting direction for future studies to determine whether adult scar-mediated healing tenocytes retain a pro-regenerative capacity that is activated upon exposure to therapeutic interventions *in vivo*.

Although healthy tenocytes were collected from young adult animals in this study, a quarter of B6 tenocytes expressed senescent ß-gal activity whereas MRL/MpJ tenocytes maintained a minimally senescent phenotype. Some studies have causally related TSDC and tenocyte senescence to pathological aging and chronic degenerative tendinopathy, whereby their accumulation leads to diminished tenogenic differentiation and cytoskeletal organization, reduced cell migration, upregulation of proinflammatory signaling cascades, and aberrant ECM remodeling (43–46). These changes extended to B6 tenocytes in our data when comparatively evaluating baseline B6 and MRL/MpJ tenocyte behavior *in vitro*. Similarly, Kallenbach and Freeberg *et al*. recently reported downregulated senescence-associated pathways in MRL/MpJ flexor tendon healing compared to that of B6 using bulk transcriptomic analyses (14). However, to our surprise and contrary to our ß-gal staining data, the static secretome of MRL/MpJ tenocytes was enriched for more known human senescence-associated secretory phenotype (SASP) factors (47) compared to that of B6 tenocytes, suggesting a potentially beneficial and tissue-dependent context for some of these SASP factors. Mounting evidence has shown that other regenerative animal models such as zebrafish and amphibians undergo tissue damage-induced cellular senescence that is localized to the injury site (48,49). Additionally, genetic or pharmacological (e.g., senolytic and senomorphic agents) removal of senescent cells *in vivo* has resulted in both reparative and impaired healing outcomes (49–51), underscoring divergent roles of developmentally- and injury-induced cellular senescence during wound healing. Considering that MRL/MpJ-derived components shifted B6 tenocytes towards a pro-regenerative phenotype without affecting the number of senescent cells, we speculate that these MRL/MpJ-enriched proteins overcome the degenerative ECM response associated with senescence and maintain some degree of phenotypic reprogramming even on senescent cells, as evidenced by consistent changes to their single-cell morphology. Future work should determine the extent to which these senescent tenocytes react to treatment by examining single-cell parameters in ß-gal-positive tenocytes via multiplexed IF staining.

Our findings establish that the enhanced cell contractility that defines the MRL/MpJ tenocyte phenotype are stabilized by the formation of cellular protrusions and focal adhesions. Scar-mediated rodent tenocytes treated with MRL/MpJ-derived components or our recombinant protein therapeutic not only mimicked these cellular features but also activated a more contractile myofibroblastic phenotype. Relatedly, Alisafaei *et al*. (52) recently discovered a positive two-way feedback loop between cellular protrusion dynamics and local ECM and mechanical anisotropy that promote fibroblastic activation and effective collagen remodeling. Moreover, we and others (53) have shown that connexin-43 localizes along tenocyte processes before assembling into cell-cell gap junctions at higher cell densities. Connexin-43 is the most abundant gap junction present in adult tendons, forming longitudinal and lateral contacts that regulate cellular metabolism, mechanosensitivity (e.g., stretch-induced calcium and ATP release), and intercellular communication through paracrine and autocrine effects (54–56). Taken together, these results suggest that early upregulation of connexin-43 activity through mitogenic stimulation with MRL/MpJ-derived components or the recombinant protein therapeutic facilitates an expedited healing response characterized by increased tenocyte proliferation, migration, and fibronectin deposition. Fibronectin is key constituent of the provisional ECM that guides cell adhesion, collagen assembly and fibrillogenesis, and temporal sequestration and release of soluble factors to further modulate cell behavior (57,58). Aberrant fibronectin accumulation is associated with long-term myofibroblast survival and persistence that ultimately drives scar-mediated tendon healing (59–61). In the context of the proliferative phase of acute tendon healing, our *in vitro* data suggest that MRL/MpJ tenocytes rapidly proliferate and undergo transient increases in αSMA expression and cell contractility, which self-regulates their cytoskeletal organization to enable early deposition of highly aligned fibronectin networks that promote cell migration and remodeling activity. Supporting these proposed cellular mechanisms, cell proliferation and apoptosis remained significantly elevated in MRL/MpJ tendons compared to B6 tendons at 1 and 4 wpi following acute injury *in vivo* in the present study. This data indicates a high turnover capacity of the MRL/MpJ tendon environment in actively recruiting cells throughout the proliferative and remodeling phases of healing to resolve the injury site while simultaneously preventing excessive ECM deposition that leads to scar formation. Overall, these findings shed light into the biological mechanisms that can inform the development of therapeutics for mammalian tendon regeneration.

This study is not without some limitations. First, the tendon sheath was excluded when isolating primary tendon cells as established by Bi *et al* (62). As this study also did not assess the presence of TSPCs *in vivo*, the precise functional contributions of sheath-derived TSPCs in MRL/MpJ tendon regeneration remain to be elucidated. Second, Quantibody-based proteomics are limited to preselected cytokine targets, are unable to distinguish between latent and active forms (e.g., matrix metalloproteinases; MMPs), and typically cannot recognize pro-peptides and membrane-bound cytokines (e.g., activin, IGF, TGFß). Despite these concerns, Quantibody assays offer pg-level sensitivity greater than that of traditional liquid chromatography-mass spectrometry (LC-MS) techniques (63) and thus allow for the quantitative detection of rare and low-abundance soluble proteins that are capable of engaging in ligand-receptor signaling. However, LC-MS technologies remain the analytical standard for deciphering a wide range of large intact proteins that comprise the ECM composition. Third, our study only analyzed male animals for generating primary cell lines, dPECM, and secretome as well as functional assays. We have recently shown that adult male and female MRL/MpJ tendons both demonstrate a regenerative capacity with vastly improved mechanical recovery compared to those of B6 mice (13). More recently, we also found that both male and female MRL/MpJ tenocytes exhibit similar or superior *in vitro* outcome measures, as detailed in the present study, compared to B6 tenocytes (64). These findings suggest that the cellular and molecular mechanisms underlying MRL/MpJ tendon regeneration are largely conserved between sexes. Future experiments should further investigate the extent of sexual dimorphism in MRL/MpJ tendon regeneration and ensure that the recombinant protein therapeutic similarly modulates female rodent tenocytes towards an MRL/MpJ-like phenotype.

There are several technical and scientific considerations in moving towards *in vivo* delivery of this recombinant protein therapeutic into clinically relevant tendon injuries in larger animal models. First, the present study only evaluated a subset of 17 out of the 29 proteins identified from the quantitative proteomic analyses of the tendon ECM and secretome. We anticipate that the incorporation of the remaining 12 proteins will amplify the observed therapeutic effects on scar-mediated tenocytes (e.g., ECM deposition). The integration of these proteins and their different molecular weights, isoelectric points, and half-life properties is non-trivial from a drug delivery perspective. Furthermore, as not all 29 recombinant proteins may be necessary to achieve therapeutic efficacy *in vivo,* other ongoing efforts will further refine this panel and assess its efficacy in tendon healing in both acute and chronic overuse injuries. Recent studies have demonstrated that functional blockade of molecular targets elevated during scar-mediated healing using neutralizing antibodies *in vivo* ameliorated overuse-induced tendinopathy in mice (65). While our study focused on evaluating the factors enriched in the MRL/MpJ tendon biological environment, it should be noted that more than half of all cytokines in the static secretome were increased or exclusively secreted by B6 tenocytes. Future studies will interrogate whether the inhibition of B6-enriched soluble factors can further augment the therapeutic potential of our MRL/MpJ-informed protein therapeutic.

We have previously demonstrated that MRL/MpJ tendons retain their improved healing capacity compared to B6 tendons when isolated from the systemic environment during *ex vivo* organ culture (12). In addition, we and other groups have shown reduced macrophage infiltration (10) and attenuated inflammatory pathways (11,13,14) in MRL/MpJ tendons compared to B6 tendons during the proliferative phase of healing. These observations motivated our overarching hypothesis that the local tissue environment is the primary driver of MRL/MpJ tendon regeneration. Interestingly, the top four soluble proteins enriched in the MRL/MpJ static secretome that were evaluated in this study – CXCL5 (LIX), CXCL1 (KC), macrophage inflammatory protein 2 (MIP-2), and semamorphin-3F (SEMA3F) – are chemokines and cytokines known to be potent mediators of early neutrophil recruitment and retainment (66–70). In contrast, decreased circulating serum levels of LIX, KC, MIP-2α during acute tendon injury have been attributed to the regeneration of MRL/MpJ ear wounds (68) and tendons (14). Therefore, we postulate that altered crosstalk between tenocytes and tissue-resident immune cells (e.g., macrophages) or locally directed molecular cues may prompt neutrophils to initiate MRL/MpJ tendon healing by concurrently streamlining the clearance of damaged tissue and the production of the provisional ECM. Nonetheless, future studies should resolve the temporal progression of the inflammatory cascade in MRL/MpJ tendon regeneration and assess the potential immunomodulatory effects of the recombinant protein therapeutic *in vivo*. Lastly, our study identified dual-specificity phosphatase 1 (DUSP1) as the most activated upstream regulator in the B6 static secretome. DUSP1 is capable of dephosphorylating major cell signaling pathways involved in innate immunity, cellular metabolism, and ECM remodeling, including extracellular-signal regulated kinase (ERK) 1/2, mitogen-activated protein kinase (MAPK), and c-Jun N-terminal kinase (JNK) (71–73). Conversely, upstream analysis of the MRL/MpJ static secretome revealed activation of ERK, p38 MAPK, and Jun, thus suggesting that these pathways may be upregulated in MRL/MpJ tendon regeneration and could be the basis for future mechanistic interrogation.

In summary, we demonstrate that the biological mechanisms underlying MRL/MpJ tenocyte behavior can be induced across different mammalian species using a subset of recombinant proteins, which may ultimately have transformative impact for promoting tendon regeneration.

## Materials and Methods

### Experimental Design

The objective of this study was to assess the utility of MRL/MpJ tendon ECM- and secretome-derived proteins in promoting regenerative outcomes across different species of tenocytes. First, we identified cellular and biological processes that delineate scar-mediated B6 and regenerative MRL/MpJ tenocyte behavior using primary cell culture assays and confocal microscopy. Next, we demonstrated that the combined administration of secretome and decellularized provisional ECM (dPECM) obtained from healthy MRL/MpJ cells and hole-punched tendons at 7 dpi, respectively, modulates B6 cells toward an MRL/MpJ-like phenotype *in vitro*. Lastly, we identified key constituents enriched in the MRL/MpJ secretome using quantitative proteomics and subsequently evaluated the functional activity of a panel of structural and soluble recombinant proteins on rat tenocytes *in vitro*. We hypothesized that selectively delivering this panel of recombinant proteins will recapitulate the effects of the MRL/MpJ-derived components on tenocyte behavior. We further hypothesized that this therapeutic will elicit interspecies utility as a novel therapeutic strategy for tendon healing. All experiments were carried out at passage three (mouse) or passage four (rat) with a minimum of three biological replicates (*N*) unless specified otherwise. For each biological replicate, three technical replicates (*n*) were included.

### Animals and patellar tendon midsubstance punch injury

All animal procedures were approved by the Cornell University Institutional Animal Care and Use Committee (IACUC). Male 16-to 18-week-old B6 and MRL/MpJ mice were bred in-house with the original dame and sire purchased from The Jackson Laboratory (B6 #000664 and MRL/MpJ #000486), unless specified otherwise, with five animals housed per cage. Male 9-to 10-month-old SD rats were purchased from Envigo (#002) with two animals housed per cage. Animals were maintained under alternating 12-hour light/dark cycles with ad libitum access to food and water.

To generate tendon dPECM, a full-thickness midsubstance injury was introduced into the left patellar tendon of male B6 and MRL/MpJ mice as described previously(12). Briefly, mice were anesthetized with isoflurane (2% volume, 0.3 L/min) and received an intraperitoneal administration of buprenorphine (1 mg/kg body weight). The fur was then completely removed from the left leg before exposing the patellar tendon through a skin incision above the knee cavity. A custom-machined stainless-steel backing (1.5-mm width, 0.5-mm thickness) coated with a polyurethane adhesive (#1867T2; McMaster-Carr) was inserted beneath the tendon before excising the tissue using a 1-mm biopsy punch. After the backing was removed, the skin incision was closed using 6-0 Prolene sutures (#8695G; Ethicon Inc.). Mice resumed cage activity and were euthanized after 1 wpi by CO_2_ administration followed by cervical dislocation. The provisional and contralateral tendons were immediately harvested, snap-frozen in liquid nitrogen, and stored at −80°C until use.

### Primary cell culture and reagents

Low-glucose Dulbecco’s modified Eagle’s medium (#10-014; Corning) and lot-selected fetal bovine serum (FBS; #35-015-CV; Corning) were used for primary cell expansion and all experiments. Mouse cells were harvested from the patellar tendons of B6 and MRL/MpJ mice as described previously(12). The tendon sheath and fat pad were removed before enzymatic digestion with a mixture of collagenase type I (2 mg/mL) and collagenase type IV (1 mg/mL) in serum-free media for up to 2 hours at 37°C on a rocking shaker. To isolate primary tenocytes, tissue digests were passed through a 70-μm strainer. Rat cells were similarly harvested from the patellar tendons of SD rats as described above except the sheath remained intact. Independent cell isolation batches consisted of patellar tendons pooled from five mice or one rat and served as biological replicates.

Cells were cultured in DMEM supplemented with 10% FBS and 1% antibiotics-antimycotics (penicillin-streptomycin-amphotericin B; PSA; Gibco) (full-serum media) at 37°C and 5% CO_2_ with media replenished every 2 days. Cells were routinely passaged at 80-90% confluency. For 2D ECM substrates, rat tail collagen I (#A1048301; Thermo Fisher Scientific) and human plasma fibronectin (#33-016-015; Thermo Fisher Scientific) were used. All cell lines employed in this study tested negative for mycoplasma contamination (#LT07-703; Lonza).

### Immunofluorescence staining and confocal microscopy of cultured tenocytes

For all immunofluorescence experiments, cells were cultured in glass-bottom 96-well plates (#655892; Greiner Bio-One). Antibody details are available in the Supplementary Materials (**Table S1**). Cells were fixed with 2% paraformaldehyde (PFA) for 10 minutes, 4% PFA for 20 minutes, washed thrice with 1X PBS, permeabilized with 0.1% Triton X-100 for 15 minutes, and blocked with 3% bovine serum albumin (BSA; in 1X PBS) for 1 hour. Primary antibodies (in 3% BSA) were incubated overnight at 4°C unless specified otherwise. Cells were then washed thrice with 1X PBS before incubating with secondary antibodies (in 3% BSA) for 1 hour at room temperature in the dark. Lastly, cells were washed thrice with 1X PBS, counterstained with DAPI (1:1000 dilution in 1X PBS; #D1306; Thermo Fisher Scientific) for 30 minutes, and finally rinsed twice with 1X PBS before imaging. For ECM staining, primary antibodies for fibronectin and collagen I were incubated prior to permeabilization as to only label cell-deposited ECM.

Images were acquired using a Zeiss LSM710 confocal microscope (Zeiss) using a 25X magnification water-immersion objective. Identical acquisition parameters were used across samples for each independent experiment. All image analyses were conducted in ImageJ (National Institutes of Health) by a blinded user.

### Tendon decellularization and preparation of dPECM coatings

Tendons harvested from B6 and MRL mice were decellularized using a detergent-free protocol inside a biosafety cabinet (BSC) as described previously (12,74). Briefly, the tissues were incubated with 50 nM of Latrunculin B (#2182-1; BioVision) in DMEM for 2 hours at 37°C at 450 RPM to depolymerize actin networks. All subsequent incubation steps were supplemented with 1X protease inhibitor cocktail (#78425; Thermo Fisher Scientific) and conducted at room temperature at 450 RPM to minimize proteolysis. The tissues were washed with deionized water for 30 minutes after Latrunculin B and hypertonic incubation steps. After exposure to Latrunculin B, the tissues were incubated in 0.6 M KCl for 2 hours, 1.0 M KI for 2 hours, and then washed in DI water for 12 hours followed by repeated KCl and KI incubation steps to induce osmolysis. Lastly, the tissues were subsequently incubated in 1X phosphate-buffered saline (PBS) containing 1 kU/mL of Pierce^TM^ Universal Nuclease (#88702; Thermo Fisher Scientific) for 2 hours followed by two consecutive 24-hour 1X PBS wash steps to clear residual cellular debris. Decellularized tissues were lyophilized for 72 hours before being snap-frozen, mechanically homogenized (Bread Ruptor 24, Omni, Inc.) using 2.8-mm ceramic beads (#19-646; Omni, Inc.), solubilized (10 mg dry weight/mL) in 1 mg/mL pepsin (#P7012; in 0.1 M HCl; Sigma-Aldrich) for 72 hours at 4°C at 150 RPM, and stored at −80°C until use. Flat-bottom 96-well plates were coated with solubilized dPECM (1 mg dry weight/mL diluted in 0.1 M AcOH) overnight at 4°C. Coatings were washed twice with 1X PBS immediately before use to remove residual acid.

To assess the contributions of the MRL/MpJ-derived components in modulating B6 cells toward an MRL/MpJ-like phenotype, the following experimental groups were evaluated for single-cell morphology and confluent mouse cell experiments: (1) B6 or MRL/MpJ cells alone, (2) cells cultured with only dPECM, (3) cells cultured with only secretome, and (4) cells cultured with both dPECM and secretome. MRL/MpJ cells treated with MRL/MpJ-derived components served as a positive control.

### Secretome preparation and quantitative proteomics

To generate the static secretome, B6 and MRL/MpJ cells were seeded at 90% confluency on Nunc^TM^ EasYFlask^TM^ culture flasks (#159910; Thermo Fisher Scientific), rinsed once with 1X DPBS (#21-030-CV; Corning), and subsequently incubated with phenol red-free, serum-free media (#11054020, Thermo Fisher Scientific) supplemented with 2X GlutaMAX^TM^ (#35050061; Thermo Fisher Scientific) for 24 hours. The conditioned medium was then collected, concentrated using ultrafiltration centrifugal devices (#88526; 3 kDa molecular weight cutoff; polyethersulfone membrane; Thermo Fisher Scientific), and stored at −80°C until use.

To investigate the effect of mechanical stimulation on the secretome composition, silicone well plates designed for the CellScale MCFX bioreactor (CellScale Biomaterials Testing) were functionalized by surface silanization as described previously (75,76). Briefly, silicone plates were plasma treated for 5 minutes followed by incubation with 10% (3-aminopropyl)triethoxysilane (APTES; #440140; Sigma-Aldrich) in ethanol at 60°C for 60 minutes. The plates were then washed thrice with 1X PBS (#21-040; Corning), incubated with 3% glutaraldehyde (#16539-60; Electron Microscopy Sciences) for 20 minutes at room temperature, and rinsed twice with 1X DPBS before covalently coating the substrates with provisional B6 and MRL/MpJ dPECM (1 mg/mL) overnight at 4°C. B6 and MRL/MpJ cells were seeded onto their respective dPECM-coated plates in full-serum media and allowed to adhere for 12 hours to reach 90% confluency. To generate the dynamic secretome, cells were then rinsed once with 1X PBS, incubated with serum-free media supplemented with 2X GlutaMAX^TM^, and immediately subjected to a homeostatic uniaxial cyclic stretch regimen (4% sinusoidal strain at 1 Hz; additional details available in Supplementary Materials) for 24 hours before collecting and concentrating the conditioned medium as described above. Streptomycin is known to block stretch-activated ion channels (77), so cells used for dynamic secretome collection were cultured in medium without PSA.

Secretome was diluted to a 2X final working concentration for all *in vitro* experiments, which was determined as the optimal concentration for promoting B6 tenocyte proliferation (data not shown), in DMEM supplemented with 1% FBS and 1% PSA (low-serum media) to limit the effect of high-abundance serum proteins. Secretome-containing media was replenished every two days from the same batch to ensure consistency between biological replicates. For quantitative proteomics (*N* = 3 per group), conditioned media was normalized to total protein concentration as determined using the bicinchoninic acid (BCA) assay (#23225; Thermo Fisher Scientific) and outsourced to RayBiotech to analyze the cytokine concentrations of the static (Quantibody^®^ Mouse Cytokine Array Q640; up to 640 targets) and dynamic secretomes (Quantibody^®^ Mouse Cytokine Array Q4000; up to 200 targets). Secretome samples were processed and analyzed in the same batch for each loading condition to minimize variation.

### Secretome proteomics bioinformatic analyses and visualization

Any detected cytokines that were not within 75% of the lower limit of detection (LOD) of the array were omitted from subsequent analyses. Cytokines with at least two uniquely detected samples that were ≥ 1.2-fold change or only detected in one genotype were regarded as differentially expressed proteins (DEPs). No comparisons for cytokines shared between the static and dynamic secretomes were conducted as these conditions were analyzed using different array technologies. Hierarchical cluster analysis was performed using the online ClustVis (78) web tool to generate heatmap expression patterns of Z-score normalized protein data for the static and dynamic secretomes. Differences between secretome conditions were visualized by clustering biological replicates using unsupervised principal components analysis (PCA). Canonical pathways, upstream regulators, and disease and functions (*P* ≤ 0.05 and | Z-score | ≥ 1.5) were performed using Ingenuity Pathway Analysis (IPA; Qiagen). Positive and negative Z-score values indicate predicted activation and inhibition, respectively, of biological functions determined by IPA. For comparative analyses in IPA, a fold change of 2.0 was assigned as a placeholder value to cytokines that were only detected in one genotype. Functional annotation of KEGG pathway enrichment analyses and GO terms were performed in R and DAVID, respectively.

### Immunohistochemical analysis of cell proliferation and apoptosis in B6 and MRL/MpJ acute tendon injuries

Male 12-to 13-week-old B6 and MRL/MpJ mice were purchased from The Jackson Laboratory and underwent a 0.75-mm full-thickness midsubstance injury. Mice were euthanized after 1 and 4 wpi to harvest the provisional tendons. Contralateral patellar tendons harvested at 24-hours post-injury from a separate study were used as naïve uninjured controls. Tendons were fixed in zinc formalin (#5701ZF, Thermo Fisher Scientific), embedded in paraffin, and sectioned coronally at a thickness of 6 µm. All subsequent immunohistochemical staining and image analysis were performed at the midsubstance injury site by a blinded user. To measure cell proliferation and apoptosis, sections were stained for cleaved caspase 3 and Ki-67, counterstained with toluidine blue, and then imaged and stitched (Zeiss AxioVision; Oberkochen, Germany) at 20X magnification. The number of positive and total cells were counted at the injury site in ImageJ to calculate the percentage of apoptotic cells (*N* = 6-9 per group). Sections were also stained for fibronectin using a custom MATLAB code to quantify the percentage of positively-stained matrix (*N* = 6-9 per group) taken at 5X magnification.

### Cell senescence and population growth assays

To evaluate the effect of serial passaging on cell growth, B6 and MRL/MpJ cells were continuously cultured from P2-P4 (*N* = 5 per group averaged from n = 3 technical replicates per *N*) and counted using a hemocytometer to calculate population doubling level (PDL; **Equation 1**) and population doubling time (PDT; **Equation 2**):

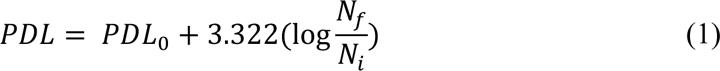

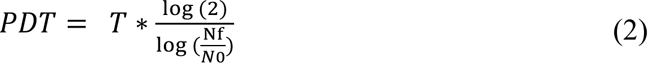

Where N_f_ is the cell number at the time of passage, N_i_ is the initial cell number seeded into the flask, PDL_0_ is the population doubling level of the previous passage, and T is the culture period in hours. B6 and MRL/MpJ cells were then seeded at P3-P5 (5,000 cells/well) onto uncoated substrates in full-serum media and allowed to adhere for 24 hours. Cells were stained for β-galactosidase (β-Gal) as a marker of senescence using a commercial kit (#9860S; Cell Signaling Technology) and imaged under an inverted brightfield microscope at 10X magnification. The percentage of senescent cells (*N* = 5 per group averaged from n = 3 technical replicates per *N*) was quantified at P3-P5 by counting the number β-gal-positive and total cells. Static secretome DEPs were mapped onto a published human SASP proteomic atlas(47) to correlate cellular senescence to biological function.

To investigate the role of the dPECM and secretome in regulating cell senescence and proliferation, B6 and MRL/MpJ cells were seeded (5,000 cells/well) onto uncoated or dPECM-coated substrates in full-serum media and allowed to adhere for 24 hours. Cells were then cultured in either low-serum or secretome-containing media for 3 days and analyzed for β-Gal staining (*N* = 5 per group averaged from n = 3 technical replicates per *N*) as described above.

### Immunofluorescence staining for p21 and DNA damage

To measure p21 activity and intrinsic DNA damage, B6 and MRL/MpJ tenocytes were seeded in full-serum media at a moderate density (5,000 cells/well) on uncoated substrates and allowed to adhere for 24 hours. Cells were then washed once with 1X DPBS and subsequently cultured in low-serum media for 24 hours prior to staining for p21 and γH2AX. The nuclear p21 integrated density was averaged for all cells in each representative image (*N* = 3 and *n* = 24 images per group). For γH2AX, the number of positive foci (*N* = 3 and *n* = 215-216 cells per group) and pan-nuclear integrated density were measured. (*N* = 3 and *n* = 24 images per group).

### Pharmacological inhibition of cytoskeletal tension and organization

B6 and MRL/MpJ cells were seeded (750 cells/well) onto collagen I-coated (50 μg/mL) substrates in full-serum media and allowed to adhere for 24 hours. To modulate cytoskeletal tension, cells were treated for 4 hours with inhibitors against myosin II (blebbistatin at 50 μM in DMSO; #B0560; Sigma-Aldrich), ROCK (Y-27632 at 50 μM in DMSO; #1254; Tocris Bioscience), and actin polymerization (Latrunculin A at 0.5 μM in DMSO; #10010630; Cayman Chemical Company). Cells treated with 0.1% DMSO served as controls.

To assess differences in B6 and MRL/MpJ cell mechanotransduction and morphology, cells were stained for the YAP and F-actin. YAP nuclear localization (nuclear-to-cytoplasmic mean intensity ratio) and single-cell morphology (circularity; **Equation 3**, perimeter, area, and F-actin integrated density) were thresholded and quantified (*n* = 72 cells per group, 8 cells per well).

### Cell migration assay

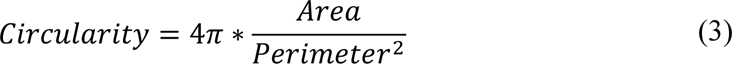

For 2D cell migration experiments (*N* = 5 per group averaged from n = 3 technical replicates per *N*), 96-well plates were coated overnight at 4°C with collagen I (50 μg/mL; #A1048301; Thermo Fischer Scientific) or fibronectin (50 μg/mL; #33-016-015; Thermo Fisher Scientific) to recapitulate biochemical cues associated with homeostasis and wound healing, respectively. B6 and MRL/MpJ cells were seeded (7,500 cells/well) onto collagen I- and fibronectin-coated substrates in full-serum media and allowed to adhere for 12 hours. Cells were serum starved in low-serum media for 18 hours to induce cell cycle synchronization, manually scratched using a 200 μL pipette tip, rinsed once with 1X DPBS, and subsequently cultured in low-serum media.

Transmitted light images were taken using a SpectraMax i3x Multi-Mode Microplate Reader (Molecular Devices). The number of cells that migrated into the defect region was quantified for 0- and 12-hours post-wounding.

### Modulating single-cell morphology, cytoskeletal protrusions, and focal adhesions

To assess whether MRL/MpJ components are capable of shifting B6 cell morphology toward previous observations(12) of decreased cell circularity and enhanced cytoskeletal protrusions, B6 and MRL/MpJ cells were seeded (750 cells/well) onto collagen I-or dPECM-coated substrates in full-serum media and allowed to adhere for 18 hours. Cells were serum starved in low-serum media for 24 hours and cultured in either low-serum or secretome-containing media for 24 hours. To assess differences in cell morphology and focal adhesion formation, cells were stained for F-actin and vinculin, respectively.

Single-cell morphology was thresholded and quantified. Cytoskeletal protrusion metrics (total number of protrusions, maximum protrusion length, total protrusion length) were manually measured using the freehand line tool for all protrusions greater than 25 µm in length. Semi-automated quantification of focal adhesions (total number of focal adhesions, average focal adhesion area, number of focal adhesions normalized to cell area) was performed as described previously (79). All experiments were analyzed with *N* = 3 per group and up to *n* = 54 cells per group with an equal number of cells randomly selected from each well.

### Stimulating cytoskeletal alignment, matrix deposition, and gap junction formation

B6 and MRL/MpJ cells were seeded (2,000 cells/well) onto uncoated substrates and cultured in full-serum media for 3 or 7 days. Media was changed every 2 days. To compare baseline behavior for cytoskeletal alignment, matrix deposition, intercellular communication, and myofibroblast activity, cells were stained for F-actin, fibronectin, collagen I, Cx-43 gap junctions, and αSMA, respectively. Cytoskeletal alignment was calculated as the local coherency/alignment index of F-actin stress fibers averaged over five elliptical regions of interest (ROIs) for each representative image using the OrientationJ (80) plugin. A coherency value of 1 indicates high anisotropy with perfectly aligned fibers, whereas a value close to 0 indicates high isotropy with no preferential fiber alignment. Cx-43 levels were measured as the integrated density normalized to cell number. Fibronectin fibril alignment was similarly analyzed as described above. Fibronectin and collagen I deposition were measured by normalizing the integrated density to the cell number of each representative image. Myofibroblast activity was measured total αSMA integrated density. Fibronectin and collagen I thickness were measured as the maximum intensity projection of Z-stack images (*N* = 3 and *n* = 27 Z-stacks per group). All other outcome measures were analyzed with *N* = 3 per group and up to *n* = 45 images per group with an equal number of images randomly taken in each well.

To assess whether MRL/MpJ components are capable of shifting confluent B6 cells toward an MRL/MpJ-like phenotype, B6 and MRL/MpJ cells were seeded (7,500 cells/well) onto uncoated or dPECM-coated substrates in full-serum media and allowed to adhere for 24 hours. Cells were then cultured in either low-serum or secretome-media for 7 days. Media was changed every 2 days. Cytoskeletal alignment, matrix deposition, intercellular communication, and myofibroblast activity were assessed by staining and analyzing cells for F-actin, fibronectin, Cx-43, and αSMA. Cells were also stained for collagen I deposition by measuring the integrated density normalized to cell number. Fibronectin fibril alignment could not be analyzed due to insufficient ECM deposition under low-serum culture conditions. All outcome measures were analyzed with *N* = 3 per group and up to *n* = 45 images per group with an equal number of images randomly taken in each well.

### Development of recombinant protein therapeutic and functional activity on rat cells

Proteomic analyses of the mouse patellar tendon (MRL/MpJ 1-week dPECM compared to B6 uninjured dECM) and secretome identified proteins of interest (POIs) for in vitro investigation. Structural POIs with at least two unique peptides and that were ≥ 2-fold higher in the MRL/MpJ 1-week dPECM (*P* ≤ 0.1) were collectively applied onto collagen I-coated substrates. Soluble POIs with at least two detected samples and that were ≥ 1.2-fold higher (*P* ≤ 0.05) were supplemented in low-serum media. A candidate panel of 29 POIs (10 structural and 19 soluble) were identified, 17 of which were evaluated as a preliminary subset in this study (**Table 1**). All POIs were prepared fresh from the same batch of stock solution to minimize the impact of freeze-thawing on protein stability. Structural POIs were administered by first applying acid soluble proteins (in 0.1 M AcOH) onto an existing base collagen I coating (50 µg/mL) overnight at 4°C. After washing twice with 1X PBS to remove residual acid, non-acid soluble POIs were then applied (in 1X PBS) overnight at 4°C. After washing twice again with 1X PBS, SD tenocytes were seeded (750 cells/well) onto POI-coated substrates in full-serum media and allowed to adhere for 18-24 hours.

To assess whether the recombinant protein therapeutic elicits a similar effect as MRL-derived components across different species of tenocytes, the following experimental groups were evaluated for single-cell morphology and confluent cell experiments: (1) rat cells alone, (2) cells cultured with dPECM (B6 uninjured or MRL/MpJ 1-week provisional) and secretome (B6 or MRL/MpJ), and (3) cells cultured with the POIs. For single-cell experiments, adhered SD tenocytes were serum starved in low-serum media for 24 hours and cultured in either low-serum or soluble POI-containing media for 24 hours. For confluent cell experiments, adhered SD tenocytes were cultured in either low-serum or soluble POI-containing media for 7 days. Media was changed every 2 days.

Rat single-cell morphology, cytoskeletal protrusion metrics, and aSMA integrated density were analyzed as described above (*N* = 3 and *n* = 72 cells per group). Cytoskeletal alignment and fibronectin deposition were analyzed as described above (*N* = 3 and *n* = 45 images per group).

### Statistical Analysis

Baseline comparisons between B6 and MRL/MpJ cells or B6 and rat cells were performed using a Welch’s *t*-test for single-cell morphology and confluent cell parameters, a Mann-Whitney *U* test for integrated density data, a Kolmogorov–Smirnov test for number of cytoskeletal protrusions, or a Student’s *t*-test for all other outcome measures. Comparisons between treatment groups were performed using a one-way Welch ANOVA with Games-Howell’s multiple comparisons test for single-cell morphology and confluent cell parameters, or a Kruskal-Wallis test with Dunn’s multiple comparisons test for integrated density data. Cell senescence and population growth kinetics were compared between mouse genotypes and passage numbers using a two-way analysis of variance (ANOVA) with Bonferroni or Sidak multiple comparisons test. The Shapiro-Wilk test was performed to test for normality. All statistical analyses were performed using GraphPad Prism 9.3.1 (GraphPad Software, La Jolla, CA) unless specified otherwise. Statistical significance was set at α = 0.05. Data are shown as mean ± standard error of the mean (SEM).

## Acknowledgments

We acknowledge K. Adebowale for assistance and insightful feedback in interpreting the single-cell actin cytoskeleton and focal adhesion data. We acknowledge C. Mendias and A. Shimpi for helpful discussions in interpreting the proteomics data. We also acknowledge Y. Wang for access to their lyophilizer. We acknowledge R. Bell and M. Pacheco for their contributions in developing the recombinant protein panel. We acknowledge E. Maloney for performing rat tenocyte isolations. We also acknowledge the Cornell Center for Animal Resources and Education (CARE) staff for animal care. All graphical schematics and illustrations were generated using BioRender.

## Author contributions

https://www.elsevier.com/authors/policies-and-guidelines/credit-author-statement

Conceptualization: JCM, NAP

Methodology: JCM, NAP

Investigation: JCM, EJL, HC, DAS

Visualization: JCM, EJL, HC, DAS

Supervision: NAP

Writing—original draft: JCM

Writing—review & editing: JCM, NAP

JCM and NAP conceived the study and developed the hypotheses. JCM designed experiments and performed all protocol development, animal surgeries, and research. JCM and EJL carried out proteomics analyses and confocal microscopy experiments. JCM, EJL, and HC analyzed immunofluorescence and staining data. DAS performed and analyzed histological data. NAP supervised the work. JCM wrote the paper. JCM and NAP edited and revised the manuscript.

## Data and materials availability

All data are available in the main text or the supplementary materials. Raw data for all Quantibody experiments will be made publicly available for access upon publication.

