## Supplemental Materials for "Proteins Derived From MRL/MpJ Tendon Provisional Extracellular Matrix and Secretome Promote Pro-Regenerative Tenocyte Behavior"

**This PDF file includes:**

Figs. S1 to S8

Tables S1 to S3

**Fig. S1**. **Validation of quantitative proteomics analyses for the static and dynamic secretomes generated by mouse tenocytes**. (**A**) Secretome protein concentrations were not different between B6 and MRL/MpJ samples for static or dynamic culture conditions (*N* = 3 per group). Significance was determined using an unpaired Student’s t-test when comparing genotypes for each secretome condition. (**B**) Top 10 upstream regulators identified from IPA revealed little agreement in Z-score activation patterns between the static and dynamic secretomes. Blue and red colors indicate regulators predicted to be activated or enriched in B6 or MRL/MpJ secretomes, respectively. (**C**) Pro-inflammatory cytokines known to be influenced by cyclic mechanical stimulation are differentially modulated between the static and dynamic secretomes in our dataset (*N* = 2-3 per group. Data is represented as the mean with individual data points for each biological replicate.

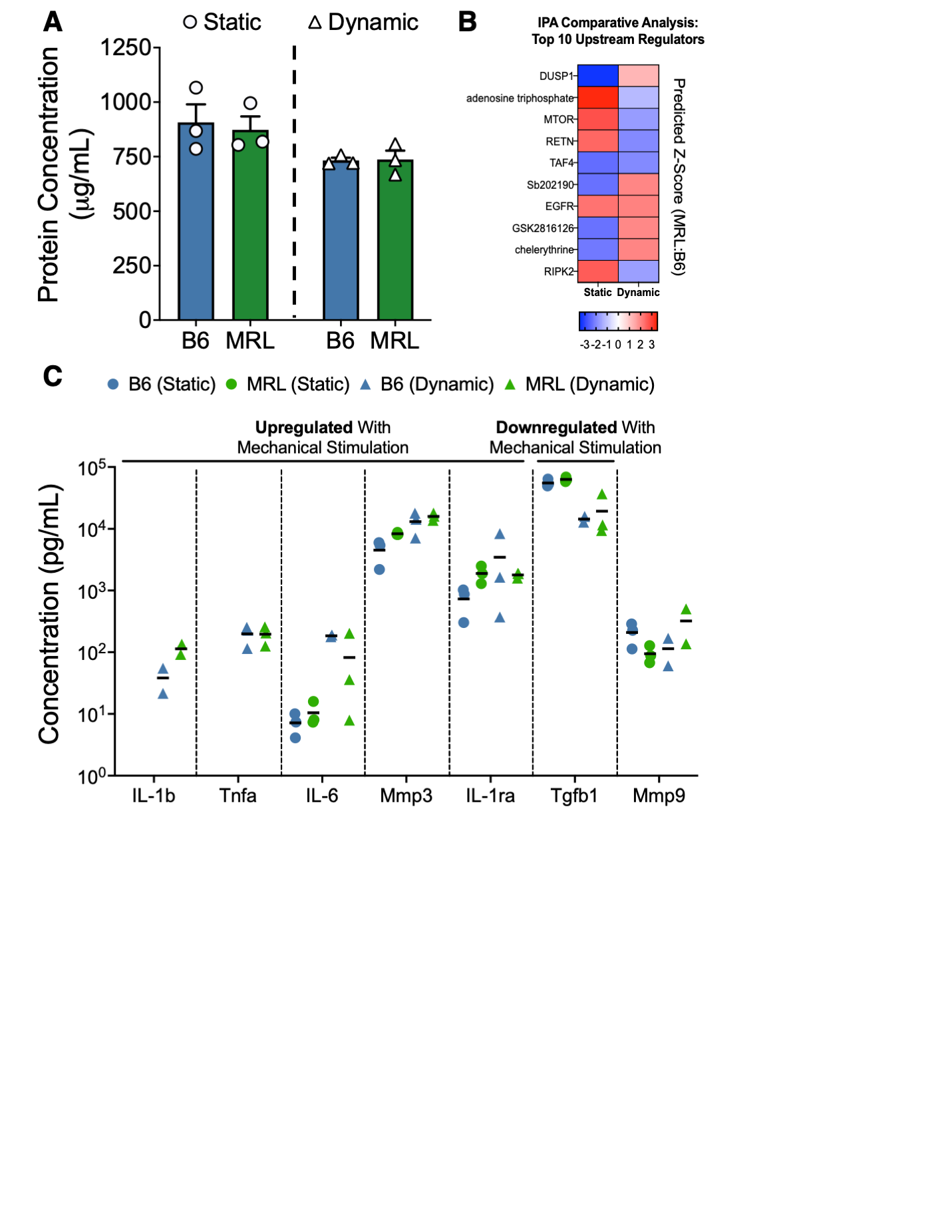

Fig. S2. Visualization of biological pathways and annotations enriched in the static secretome generated by mouse tenocytes. (A) Dot plot representation of the top 30 biological pathways (*P* < 0.05) identified using KEGG overrepresentation analysis of all differentially expressed proteins (DEPs) in the static secretome. The dot size represents the gene/protein ratio whereas the color indicates the adjusted *P* value determined by Fisher’s exact test. (B and C) IPA graphical summary overviewing all biological activities and their inferred relationships within the static secretome (B). All entities met the selection criteria of *P* < 0.05 and/or activation Z-score ≥ 2 for upstream regulators and diseases and functions. Orange and blue nodes indicate activated (enriched in MRL/MpJ) and inhibited (enriched in B6) annotations, respectively. (C) IPA legend accompanying the graphical summary.

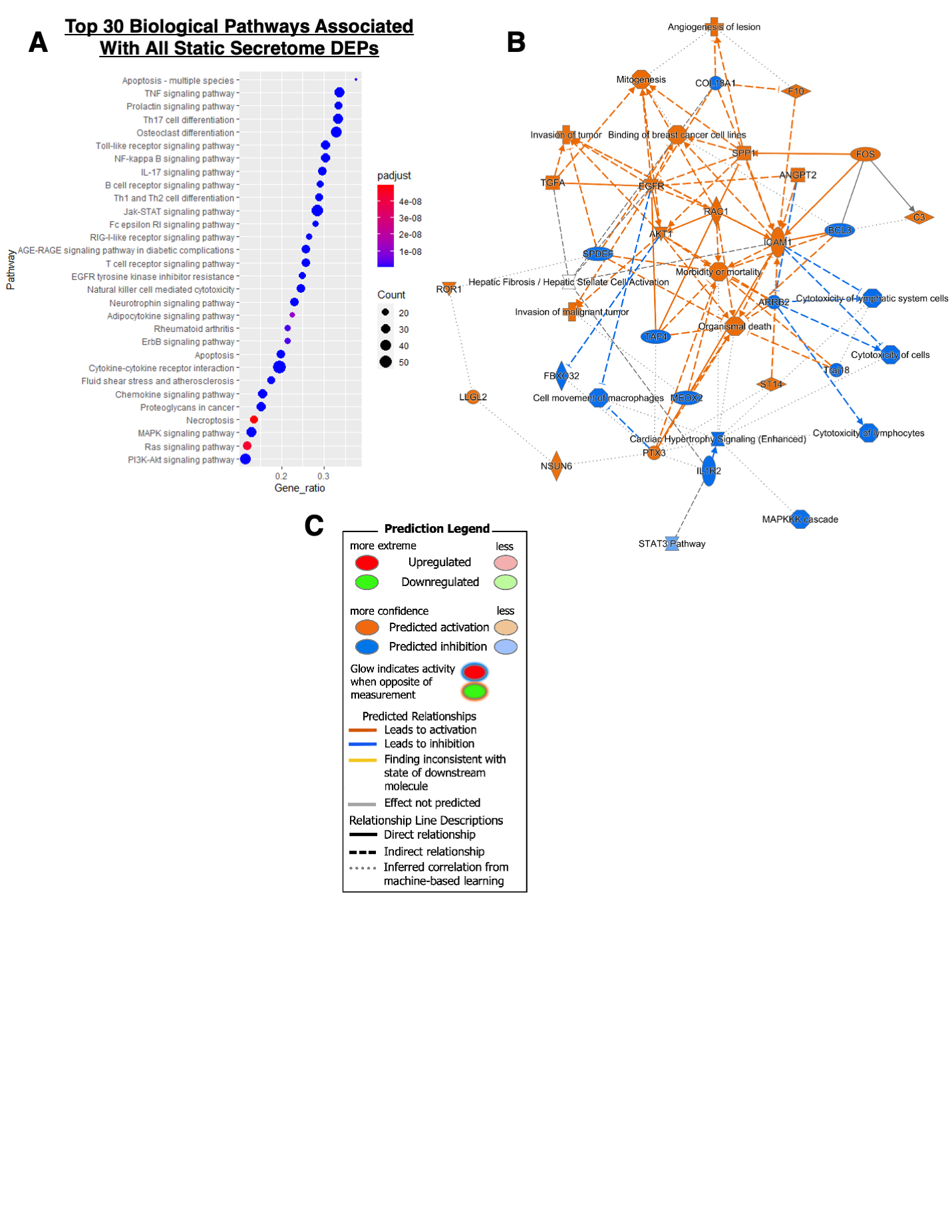

Fig. S3. B6 tenocytes progressively undergo cellular senescence with serial passages *in vitro* while total cell number remained consistent. (A) Total cell number was not different between B6 and MRL/MpJ tenocytes at each passage number or across different passage numbers for each genotype. (B) The number of senescent/ß-gal-positive B6 tenocytes significantly increased with higher passage numbers whereas MRL/MpJ tenocytes maintained a minimally senescent phenotype. **P* < 0.05, ****P* < 0.001, *****P* < 0.0001. *N* = 5 biological replicates per group averaged from *n* = 3 technical replicates per *N*. Significance was determined using a two-way ANOVA with post-hoc Bonferroni.

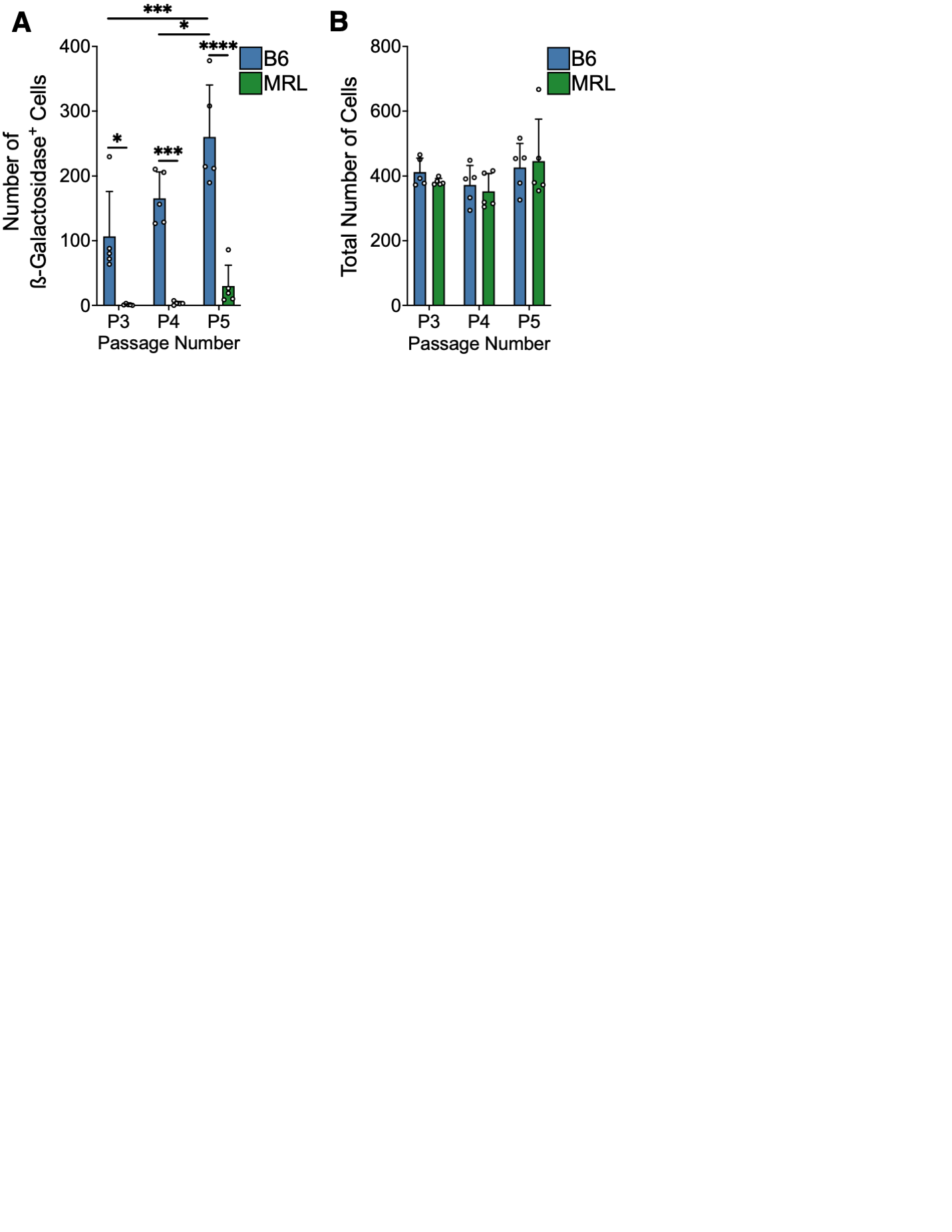

Fig. S4. B6 and MRL/MpJ tenocytes exhibit comparable intrinsic p21 expression and DNA damage profiles. (A and B) Quantification of γH2AX-positive foci (indicative of DNA damage breaks) was comparable between B6 and MRL/MpJ tenocytes (*N* = 3 with *n* = 219-220 total cells per group). (B) Pan-nuclear γH2AX fluorescence intensity (normalized to B6 mean) was significantly higher in B6 tenocytes than in MRL/MpJ tenocytes (*N* = 3 with *n* = 8 wells per group). (C) Quantification of p21 fluorescence intensity (normalized to B6 mean) was comparable between B6 and MRL/MpJ tenocytes. ****P* < 0.001. Significance was determined using a Mann-Whitney *U* test.

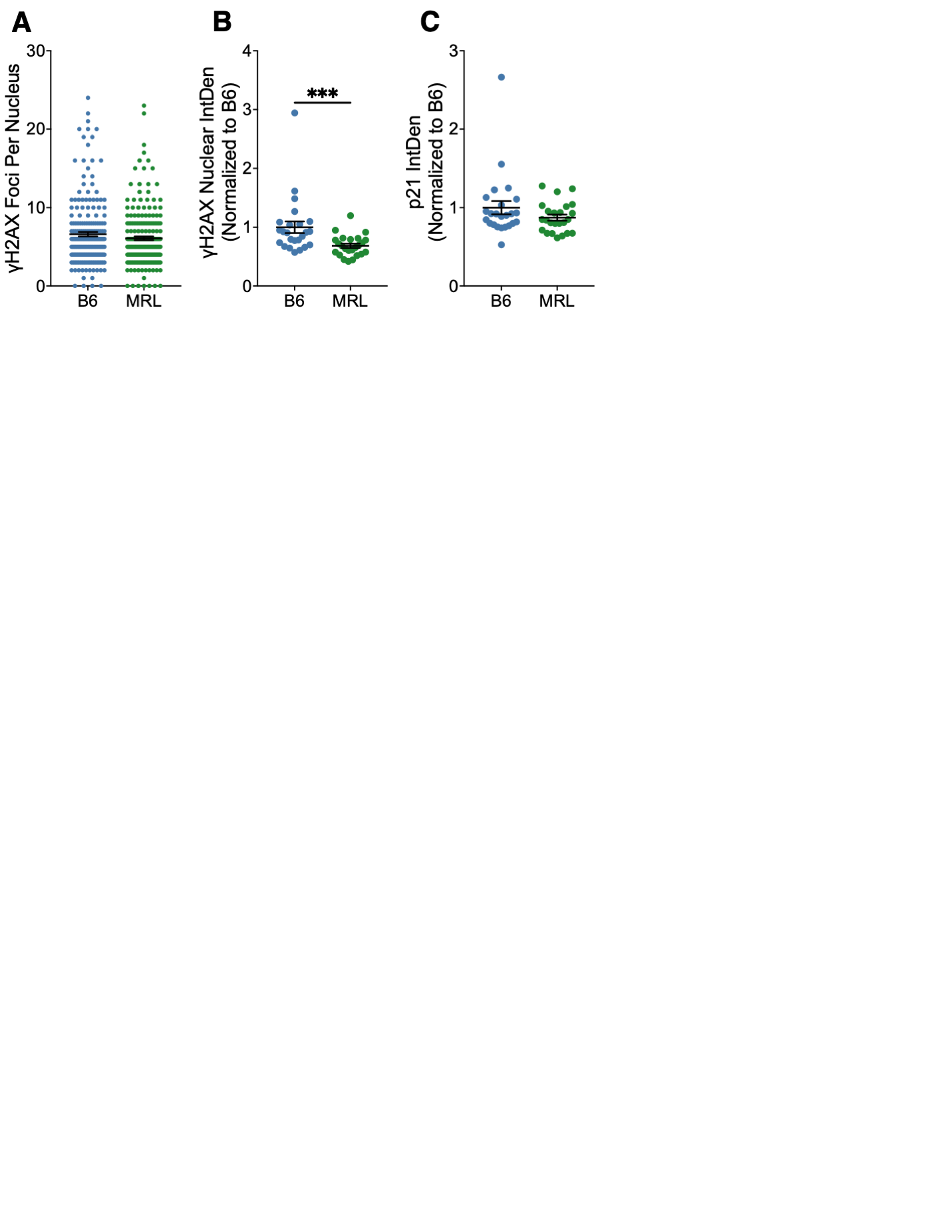

Fig. S5. Senescence-associated secretory phenotype (SASP) factors mapped to differentially expressed proteins in the static secretome are not more abundant in B6 tenocytes despite higher ß-galactosidase activity. (A) Heat map visualizing DEPs enriched in the B6 (indicated in blue; 18 total) and MRL/MpJ (indicated in red; 22 total) static secretome that were detected in a published human SASP atlas. (B and C) Breakdown of DEPs in the B6 (B) and MRL/MpJ (C) static secretomes and their contributions to the published SASP atlases of human fibroblasts (referred to as ‘fibro’), epithelial cells (referred to as ‘epi’), and exosome cargo. Warm and cool colors indicate upregulation and downregulation in the SASP, respectively. Different SASP conditions are shown (IR = x-irradiated, RAS = inducible RAS overexpression, and ATV = atazanavir treatment).

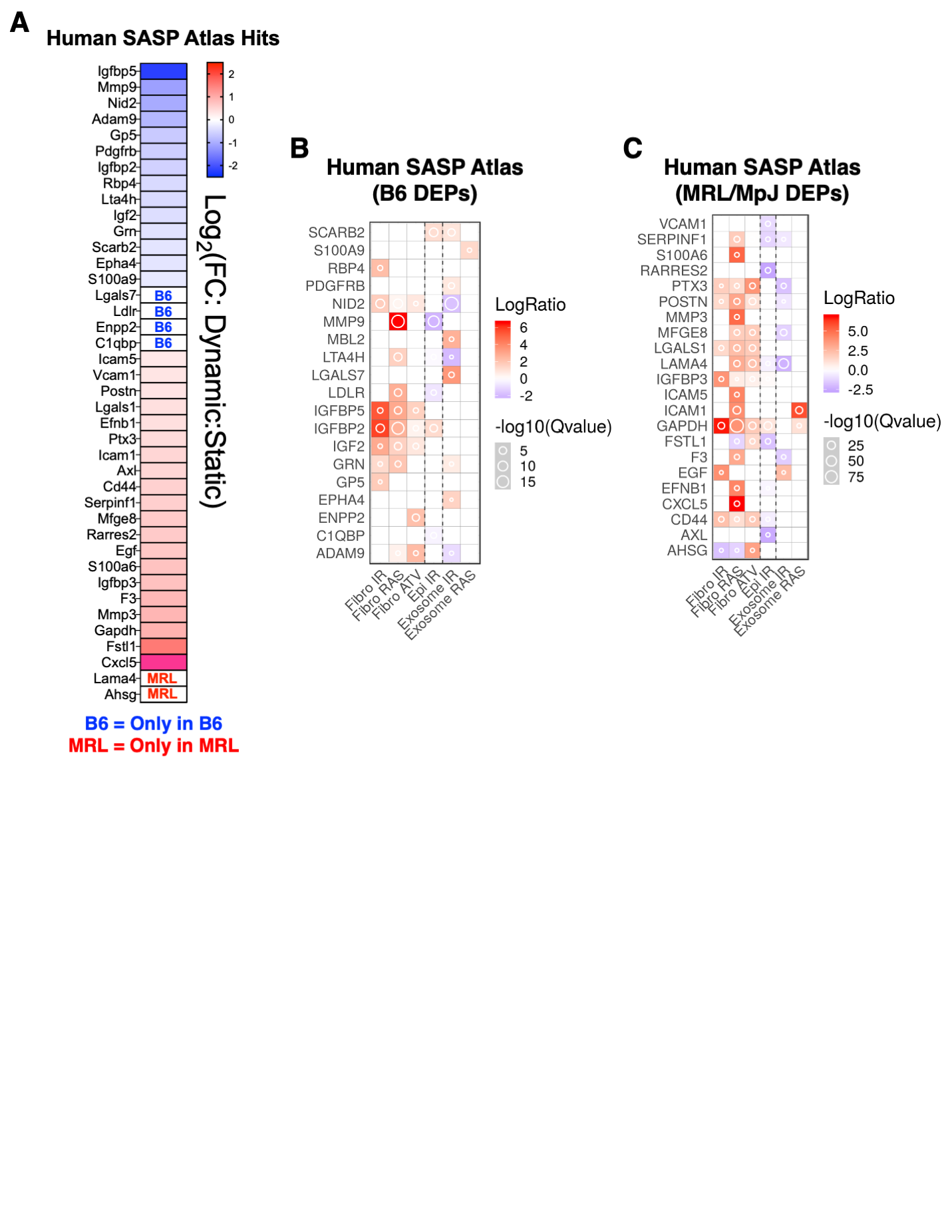

Fig. S6. MRL/MpJ-derived components increases collagen I deposition by confluent B6 tenocytes but has no effect on αSMA expression. (A) IF staining of collagen I (orange) in B6 tenocytes treated with only MRL/MpJ provisional dECM, only MRL/MpJ secretome, or a combination of mouse-derived components for 7 days (scale bar, 100 μm). (B) B6 tenocytes treated with MRL/MpJ-derived components deposited more collagen I compared to those treated with B6-derived components. (C) αSMA expression was comparable between all experimental groups. *N* = 3 with *n* = 45 representative images per group. Significance was determined using a Kruskal-Wallis test with post-hoc Dunn’s. Representative images shown for all staining experiments.

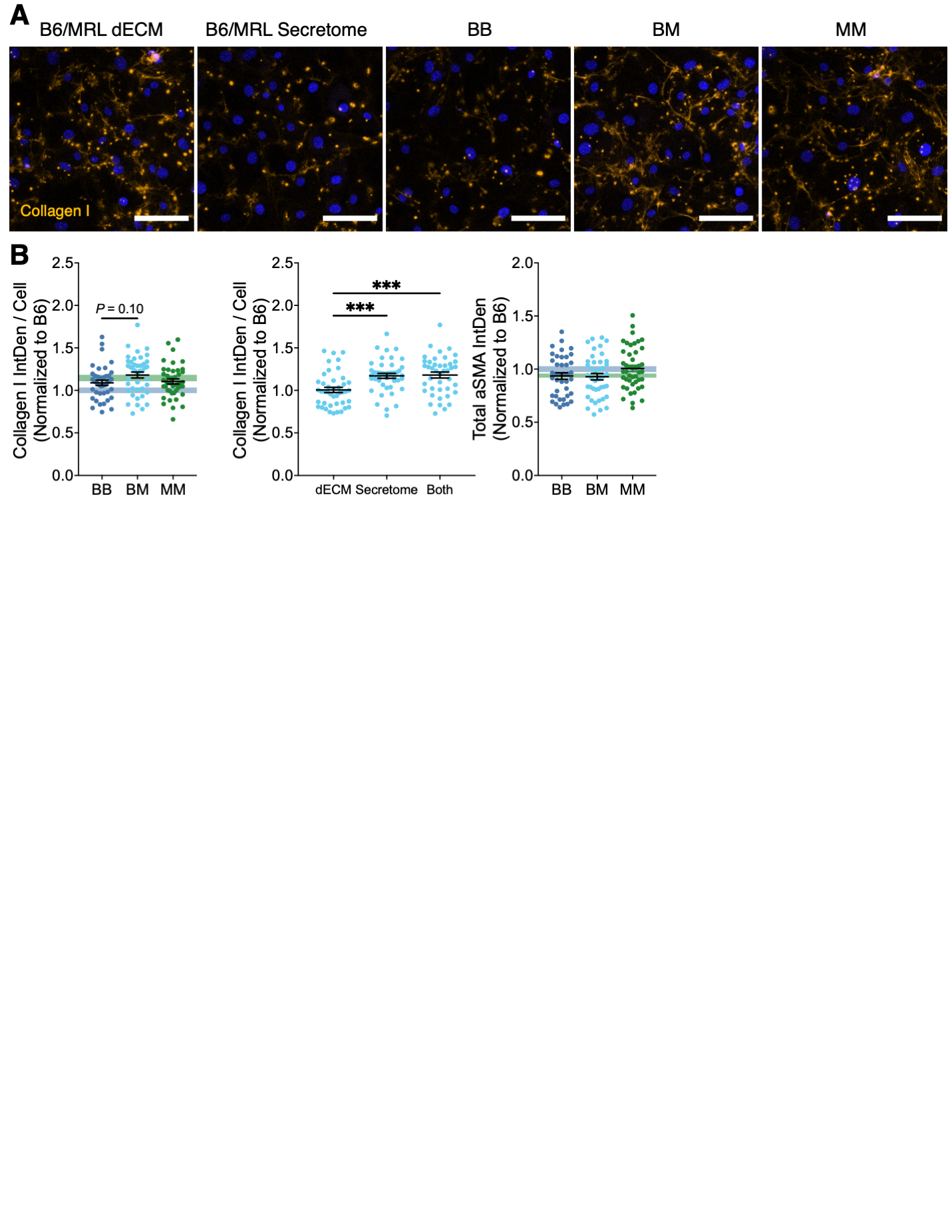

Fig. S7. Cell morphology is comparable between B6 and Sprague-Dawley (SD) tenocytes. (A and B) B6 and SD tenocytes cultured on collagen I-coated substrates displayed identical cellular protrusion characteristics (A) and single-cell morphological parameters (B). Significance was determined using a one-way ANOVA with post-hoc Tukey (percentage of cells with protrusions), Kruskal-Wallis test with post-hoc Dunn’s (number of protrusions, normalized total protrusion length, aSMA expression), or a one-way Welch’s ANOVA (total protrusion length, cell circularity, spreading area, and perimeter).

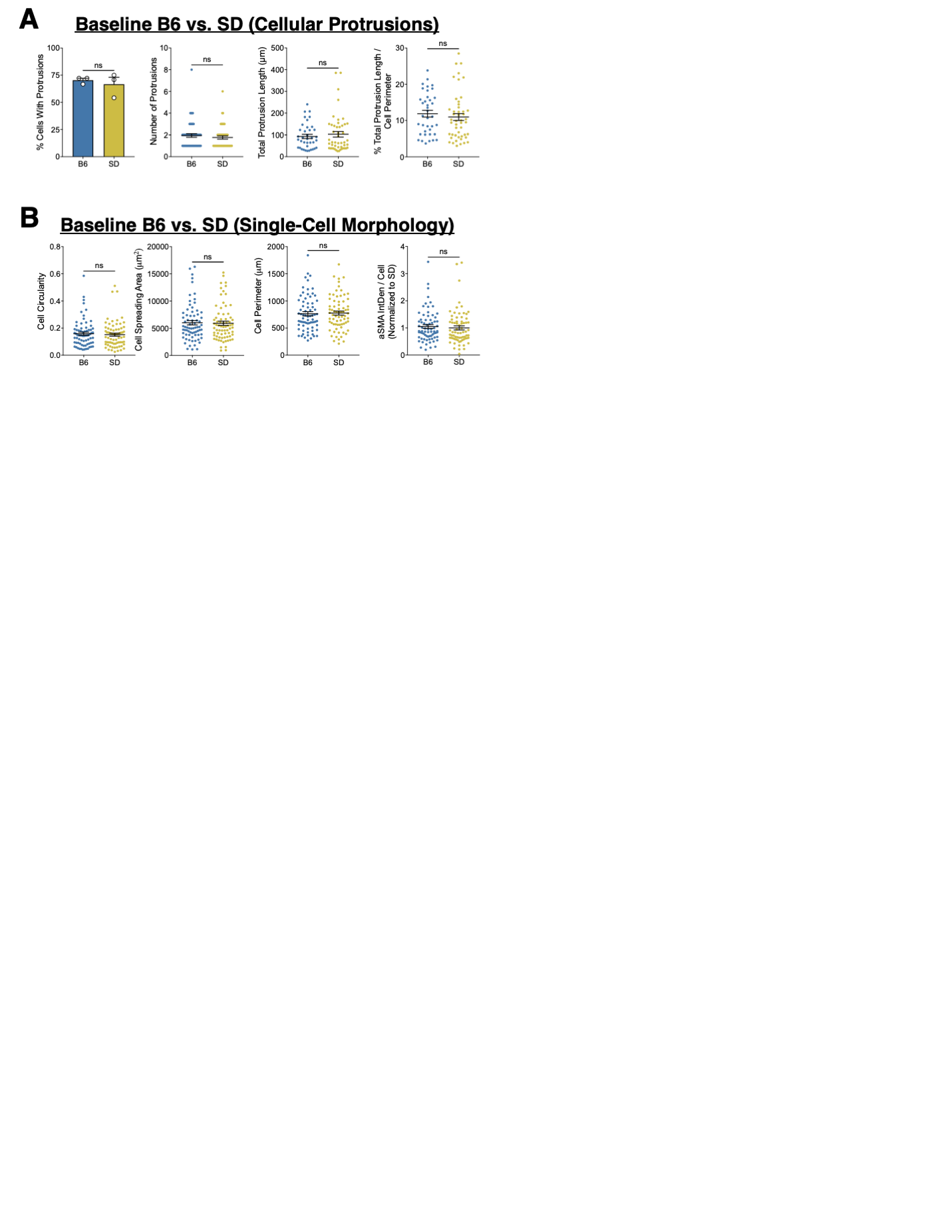

**Fig. S8. Protein concentrations of MRL/MpJ-derived cytokines from the static secretome used in the protein therapeutic**. Superimposed scatter plot illustrating fold change differences for 12 DEPs enriched in the MRL/MpJ static secretome that were included in the protein therapeutic used in this study. Data is represented as the mean withs individual data points for each biological replicate (*N* = 2-3 per group).

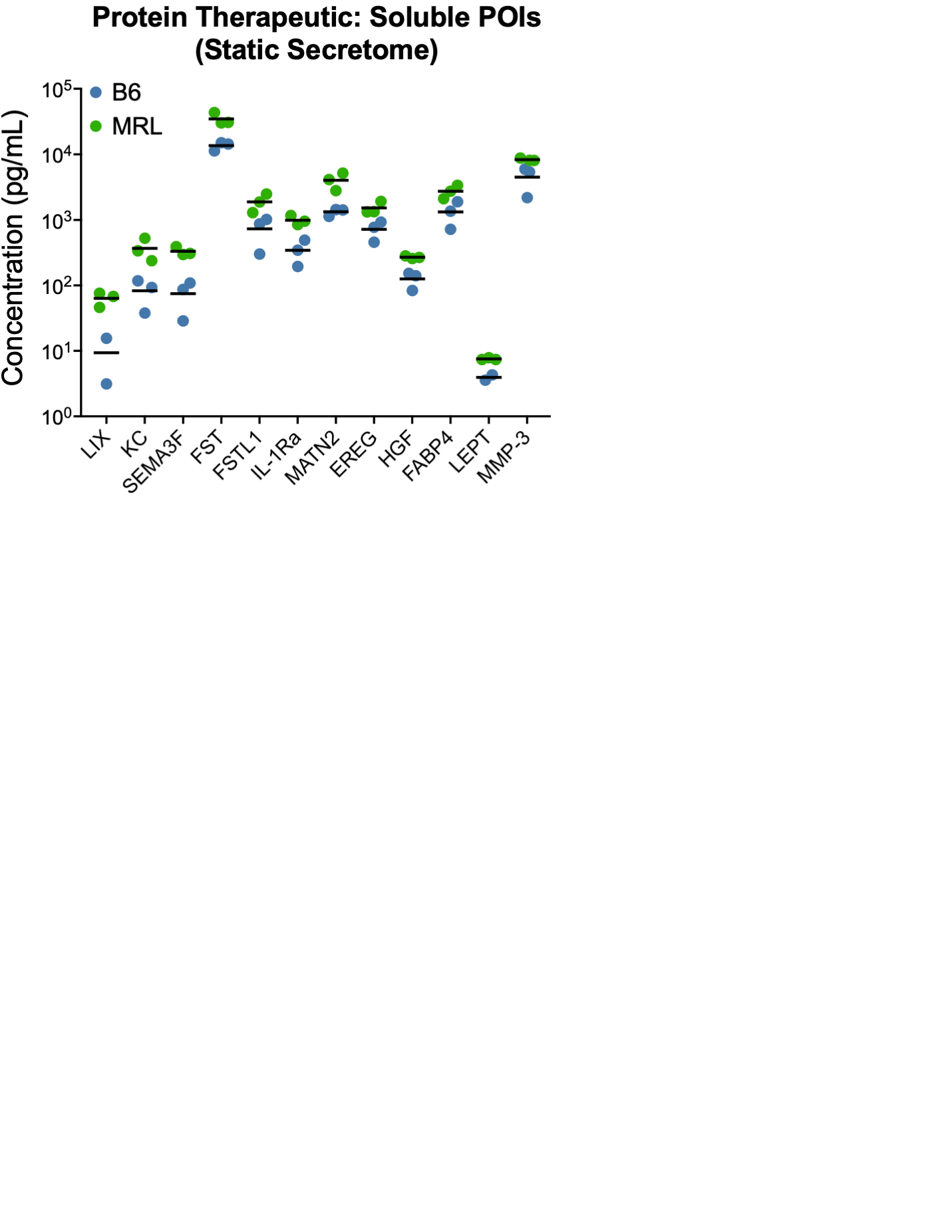

Table S1. List of primary antibodies used for immunofluorescence and immunohistochemistry

| **Antibody** | **Manufacturer** | **Catalog Number** | **Dilution and Use** |
| --- | --- | --- | --- |
| Alexa Fluor^TM^ Plus 555 Phalloidin | Thermo Fisher Scientific | A30106 | 1:400, 1 hour at room temperature |
| Vinculin | Sigma-Aldrich | V9131 | 1:400, overnight at 4°C |
| YAP | Santa Cruz Biotechnology, Inc. | sc-101199 | 1:200, 2 hours at room temperature |
| Fibronectin | Abcam | ab45688 | 1:400, overnight at 4°C |
| Cx-43 | Abcam | ab11370 | 1:1000, overnight at 4°C |
| αSMA | Abcam | ab124964 | 1:200, overnight at 4°C |
| Collagen I | Abcam | ab260043 | 1:250, overnight at 4°C |
| Sox2 | Abcam | ab92494 | 1:100, overnight at 4°C |
| Nanog | Abcam | ab80892 | 1:100, overnight at 4°C |
| Oct-3/4 | Santa Cruz Biotechnology, Inc. | sc-5279 | 1:150, overnight at 4°C |
| HIF-1α | Abcam | ab179483 | 1:500, overnight at 4°C |
| p21 | Abcam | ab188224 | 1:500, overnight at 4°C |
| γH2AX | MilliporeSigma | 05-636-I | 1:500, overnight at 4°C |
| Ki-67 | Abcam | ab15580 | 1:100, 2 hours at room temperature |
| Cleaved Caspase 3 | Cell Signaling Technology | 9661S | 1:1000, 2 hours at room temperature |

Table S2. Full list of all activated and inhibited ingenuity pathway analysis upstream regulators from proteomic analyses of the static secretome

| **Upstream Regulator** | **Molecule Type** | **Predicated Activation State** | **Activation Z-Score** | ***P* Value of Overlap** |
| --- | --- | --- | --- | --- |
| **Activated in MRL/MpJ Secretome (Inhibited in B6 Secretome) – 87 Total** | | | | |
| ATP | chemical - endogenous mammalian | Activated | 3.379 | 2.13E-11 |
| FOS | transcription regulator | Activated | 3.2 | 2.66E-21 |
| ANGPT2 | growth factor | Activated | 3.086 | 1.04E-10 |
| CG | complex | Activated | 3.059 | 1.71E-17 |
| cigarette smoke | chemical toxicant | Activated | 3.034 | 2.35E-09 |
| RORC | ligand-dependent nuclear receptor | Activated | 2.8 | 4.95E-06 |
| D-fructose | chemical - endogenous mammalian | Activated | 2.791 | 2.21E-08 |
| MTOR | kinase | Activated | 2.758 | 8.07E-06 |
| 5-hydroxytryptamine | chemical - endogenous mammalian | Activated | 2.722 | 3.73E-04 |
| RAC1 | enzyme | Activated | 2.674 | 2.34E-09 |
| TGFA | growth factor | Activated | 2.622 | 3.20E-09 |
| OLR1 | transmembrane receptor | Activated | 2.607 | 1.80E-08 |
| S-nitroso-N-acetyl-DL-penicillamine | chemical reagent | Activated | 2.596 | 2.91E-06 |
| RIPK2 | kinase | Activated | 2.596 | 9.74E-08 |
| JUN | transcription regulator | Activated | 2.574 | 4.21E-25 |
| ERK | group | Activated | 2.56 | 3.21E-12 |
| HRAS | enzyme | Activated | 2.528 | 1.15E-15 |
| IL10RA | transmembrane receptor | Activated | 2.475 | 1.19E-19 |
| dextran sulfate | chemical drug | Activated | 2.448 | 1.14E-25 |
| RETN | other | Activated | 2.446 | 4.29E-09 |
| F10 | peptidase | Activated | 2.429 | 2.84E-08 |
| PTPN1 | phosphatase | Activated | 2.414 | 1.49E-05 |
| UTP | chemical - endogenous mammalian | Activated | 2.411 | 2.38E-06 |
| EPHX2 | enzyme | Activated | 2.4 | 7.39E-08 |
| collagenase | group | Activated | 2.36 | 6.75E-10 |
| Collagen type II | complex | Activated | 2.357 | 1.35E-12 |
| cisplatin | chemical drug | Activated | 2.353 | 5.26E-12 |
| 5-O-mycolyl-beta-araf-(1->2)-5-O-mycolyl-alpha-araf-(1->1')-glycerol | chemical - endogenous non-mammalian | Activated | 2.339 | 4.95E-06 |
| NFKBIB | transcription regulator | Activated | 2.306 | 1.93E-11 |
| PTX3 | other | Activated | 2.304 | 1.31E-08 |
| ICAM1 | transmembrane receptor | Activated | 2.297 | 1.17E-12 |
| IL17F | cytokine | Activated | 2.289 | 9.86E-13 |
| AKT1 | kinase | Activated | 2.288 | 3.06E-23 |
| EGFR | kinase | Activated | 2.261 | 7.24E-15 |
| thioacetamide | chemical toxicant | Activated | 2.255 | 5.27E-12 |
| MVP | other | Activated | 2.223 | 2.84E-08 |
| ROR1 | kinase | Activated | 2.219 | 1.03E-04 |
| PADI2 | enzyme | Activated | 2.219 | 5.21E-06 |
| aldosterone | chemical - endogenous mammalian | Activated | 2.215 | 4.55E-15 |
| PLCE1 | enzyme | Activated | 2.213 | 1.28E-06 |
| isobutylmethylxanthine | chemical toxicant | Activated | 2.208 | 1.81E-07 |
| ST14 | peptidase | Activated | 2.207 | 1.28E-06 |
| NfkB1-RelA | complex | Activated | 2.207 | 1.19E-14 |
| Iga | complex | Activated | 2.206 | 4.70E-07 |
| nitroarginine | chemical reagent | Activated | 2.2 | 6.31E-05 |
| Focal adhesion kinase | group | Activated | 2.197 | 5.63E-06 |
| Usp17la (includes others) | peptidase | Activated | 2.193 | 9.20E-06 |
| S1PR2 | G-protein coupled receptor | Activated | 2.188 | 5.21E-06 |
| TRAF3IP2 | enzyme | Activated | 2.184 | 9.05E-25 |
| FGFR1 | kinase | Activated | 2.183 | 4.02E-07 |
| 3-deazaneplanocin | chemical drug | Activated | 2.179 | 2.65E-03 |
| CRP | other | Activated | 2.177 | 4.01E-05 |
| TXNIP | other | Activated | 2.176 | 5.34E-04 |
| ozone | chemical toxicant | Activated | 2.176 | 4.40E-05 |
| EGR1 | transcription regulator | Activated | 2.175 | 9.48E-19 |
| Fibrinogen | complex | Activated | 2.173 | 4.70E-07 |
| Fc gamma receptor | group | Activated | 2.163 | 6.84E-06 |
| TAC1 | other | Activated | 2.161 | 7.61E-07 |
| Pro-inflammatory Cytokine | group | Activated | 2.158 | 3.13E-11 |
| NR5A2 | ligand-dependent nuclear receptor | Activated | 2.157 | 5.92E-06 |
| E. coli B4 lipopolysaccharide | chemical toxicant | Activated | 2.156 | 6.39E-20 |
| ITGB1 | transmembrane receptor | Activated | 2.126 | 3.11E-14 |
| Lh | complex | Activated | 2.123 | 4.59E-03 |
| gentamicin C | chemical drug | Activated | 2.121 | 4.79E-06 |
| CHD1 | enzyme | Activated | 2.121 | 4.84E-11 |
| 2-bromoethylamine | chemical reagent | Activated | 2.121 | 5.92E-07 |
| TIRAP | other | Activated | 2.111 | 2.28E-11 |
| PDX1 | transcription regulator | Activated | 2.111 | 1.38E-06 |
| hyaluronic acid | chemical - endogenous mammalian | Activated | 2.103 | 9.63E-20 |
| NfkB-RelA | complex | Activated | 2.102 | 5.85E-08 |
| MAP3K1 | kinase | Activated | 2.085 | 1.28E-06 |
| P38 MAPK | group | Activated | 2.081 | 1.37E-21 |
| SPP1 | cytokine | Activated | 2.063 | 4.06E-19 |
| LGALS3 | other | Activated | 2.062 | 3.31E-15 |
| PDGF BB | complex | Activated | 2.018 | 3.62E-16 |
| C3 | peptidase | Activated | 2.006 | 7.08E-14 |
| TLR7/8 | group | Activated | 2 | 3.18E-03 |
| THBS4 | other | Activated | 2 | 1.11E-03 |
| ROCK1 | kinase | Activated | 2 | 8.57E-04 |
| NSUN6 | enzyme | Activated | 2 | 2.64E-03 |
| NFAT (complex) | complex | Activated | 2 | 2.40E-05 |
| LLGL2 | other | Activated | 2 | 9.77E-04 |
| lipoarabinomannan | chemical - endogenous non-mammalian | Activated | 2 | 3.43E-04 |
| Ikb | group | Activated | 2 | 4.07E-04 |
| D-sphingosine | chemical - endogenous mammalian | Activated | 2 | 2.18E-05 |
| CXCL1 | cytokine | Activated | 2 | 2.18E-05 |
| CDKN2B-AS1 | other | Activated | 2 | 4.31E-05 |
| **Inhibited in MRL/MpJ Secretome (Activated in B6 Secretome) – 84 Total** | | | | |
| DUSP1 | phosphatase | Inhibited | -3.348 | 1.24E-13 |
| vitamin E | chemical drug | Inhibited | -3.265 | 4.17E-09 |
| beta-estradiol | chemical - endogenous mammalian | Inhibited | -3.128 | 4.84E-28 |
| SB-431542 | chemical reagent | Inhibited | -2.884 | 5.93E-11 |
| UCP1 | transporter | Inhibited | -2.813 | 3.28E-03 |
| estrogen receptor | group | Inhibited | -2.782 | 1.51E-14 |
| MEOX2 | transcription regulator | Inhibited | -2.722 | 7.88E-12 |
| ESR1 | ligand-dependent nuclear receptor | Inhibited | -2.708 | 5.43E-13 |
| NOTCH2 | transcription regulator | Inhibited | -2.588 | 5.77E-06 |
| genistein | chemical drug | Inhibited | -2.568 | 4.45E-16 |
| STK11 | kinase | Inhibited | -2.53 | 4.81E-10 |
| INPP5D | phosphatase | Inhibited | -2.456 | 1.56E-06 |
| pimagedine | chemical drug | Inhibited | -2.449 | 1.87E-06 |
| MSC | transcription regulator | Inhibited | -2.449 | 2.00E-04 |
| FBXO32 | enzyme | Inhibited | -2.438 | 1.60E-03 |
| diphenyleneiodonium | chemical reagent | Inhibited | -2.428 | 4.09E-08 |
| actinomycin D | biologic drug | Inhibited | -2.421 | 3.18E-16 |
| TAF4 | transcription regulator | Inhibited | -2.412 | 1.78E-08 |
| Traj18 | other | Inhibited | -2.401 | 4.54E-09 |
| CHRNA7 | transmembrane receptor | Inhibited | -2.397 | 9.90E-06 |
| tosylphenylalanyl chloromethyl ketone | chemical - protease inhibitor | Inhibited | -2.395 | 1.46E-06 |
| SB203580 | chemical drug | Inhibited | -2.388 | 6.20E-24 |
| Y 27632 | chemical drug | Inhibited | -2.384 | 9.29E-11 |
| metformin | chemical drug | Inhibited | -2.375 | 3.08E-08 |
| Hbb-b1 | transporter | Inhibited | -2.359 | 1.04E-05 |
| TET2 | enzyme | Inhibited | -2.353 | 2.87E-04 |
| tempol | chemical drug | Inhibited | -2.345 | 3.63E-09 |
| Sb202190 | chemical drug | Inhibited | -2.342 | 1.02E-07 |
| APC | enzyme | Inhibited | -2.338 | 1.79E-14 |
| caffeic acid phenethyl ester | chemical drug | Inhibited | -2.336 | 8.03E-10 |
| GSK2816126 | chemical drug | Inhibited | -2.328 | 1.26E-13 |
| simvastatin | chemical drug | Inhibited | -2.327 | 1.87E-15 |
| BMP4 | growth factor | Inhibited | -2.297 | 7.53E-09 |
| CCN5 | growth factor | Inhibited | -2.259 | 4.29E-09 |
| fingolimod | chemical drug | Inhibited | -2.253 | 2.77E-09 |
| STEAP3 | transporter | Inhibited | -2.236 | 1.46E-03 |
| elaidic acid | chemical - endogenous mammalian | Inhibited | -2.236 | 1.28E-02 |
| dimethyl sulfoxide | chemical drug | Inhibited | -2.236 | 4.54E-02 |
| chelerythrine | chemical drug | Inhibited | -2.236 | 2.40E-05 |
| NR1H4f | ligand-dependent nuclear receptor | Inhibited | -2.231 | 4.59E-04 |
| plerixafor | chemical drug | Inhibited | -2.219 | 1.56E-06 |
| lipoxin A4 | chemical - endogenous mammalian | Inhibited | -2.219 | 2.10E-04 |
| alpha-tocopherol | chemical drug | Inhibited | -2.219 | 3.63E-05 |
| BAPTA-AM | chemical reagent | Inhibited | -2.217 | 6.96E-07 |
| HLX | transcription regulator | Inhibited | -2.216 | 1.20E-04 |
| miR-155-5p (miRNAs w/seed UAAUGCU) | mature microRNA | Inhibited | -2.209 | 8.53E-04 |
| ANGPT1 | growth factor | Inhibited | -2.208 | 5.22E-05 |
| ARRB2 | other | Inhibited | -2.2 | 2.80E-09 |
| U73122 | chemical reagent | Inhibited | -2.195 | 4.81E-04 |
| CAT | enzyme | Inhibited | -2.195 | 1.35E-03 |
| PECAM1 | other | Inhibited | -2.194 | 1.28E-06 |
| SMAD7 | transcription regulator | Inhibited | -2.186 | 9.81E-18 |
| RCAN1 | other | Inhibited | -2.186 | 3.63E-05 |
| methotrexate | chemical drug | Inhibited | -2.184 | 1.26E-10 |
| AIRE | transcription regulator | Inhibited | -2.184 | 3.90E-07 |
| teriflunomide | chemical drug | Inhibited | -2.182 | 3.06E-07 |
| SFTPA1 | transporter | Inhibited | -2.174 | 1.77E-04 |
| IL1R2 | transmembrane receptor | Inhibited | -2.173 | 3.06E-07 |
| NLRP12 | other | Inhibited | -2.149 | 4.70E-08 |
| estriol | chemical - endogenous mammalian | Inhibited | -2.138 | 2.98E-07 |
| paricalcitol | chemical drug | Inhibited | -2.135 | 9.20E-08 |
| SPI1 | transcription regulator | Inhibited | -2.101 | 1.17E-06 |
| PD98059 | chemical - kinase inhibitor | Inhibited | -2.097 | 1.36E-26 |
| pyrrolidine dithiocarbamate | chemical reagent | Inhibited | -2.09 | 3.31E-15 |
| BCL3 | transcription regulator | Inhibited | -2.073 | 2.11E-09 |
| RARA | ligand-dependent nuclear receptor | Inhibited | -2.043 | 2.63E-06 |
| COL18A1 | other | Inhibited | -2.043 | 7.19E-10 |
| ADA | enzyme | Inhibited | -2.039 | 1.75E-10 |
| APOA1 | transporter | Inhibited | -2.027 | 2.40E-05 |
| thalidomide | chemical drug | Inhibited | -2.026 | 3.63E-09 |
| fasudil | chemical drug | Inhibited | -2.008 | 2.06E-07 |
| valsartan | chemical drug | Inhibited | -2 | 1.17E-14 |
| SPDEF | transcription regulator | Inhibited | -2 | 3.68E-03 |
| Sod | group | Inhibited | -2 | 9.89E-05 |
| SELENOS | other | Inhibited | -2 | 5.81E-05 |
| RXRB | ligand-dependent nuclear receptor | Inhibited | -2 | 3.39E-12 |
| NANOG | transcription regulator | Inhibited | -2 | 1.29E-02 |
| JMF3086 | chemical reagent | Inhibited | -2 | 5.75E-06 |
| FOXP1 | transcription regulator | Inhibited | -2 | 5.22E-05 |
| fluoxetine | chemical drug | Inhibited | -2 | 3.69E-02 |
| EPHB6 | kinase | Inhibited | -2 | 5.22E-03 |
| CASP3 | peptidase | Inhibited | -2 | 9.77E-04 |
| arginine | chemical - endogenous mammalian | Inhibited | -2 | 1.96E-03 |
| A-Fos | chemical reagent | Inhibited | -2 | 2.37E-04 |

**Table S3. Comparative ingenuity pathway analysis diseases and functions of the static and dynamic secretomes**

| **Canonical Pathways** | **Static Activation Z-Score** | **Static *P* Value of Overlap** | **Dynamic Activation Z-Score** | **Dynamic *P* Value of Overlap** |
| --- | --- | --- | --- | --- |
| Aggregation of cells | -1.967 | 2.51E-05 | -2.295 |  |
| Cell movement of macrophages | -2.03 |  | -2.191 |  |
| Organismal death | 2.286 |  | 1.905 |  |
| Morbidity or mortality | 2.203 |  | 1.975 |  |
| Angiogenesis of lesion | 2.459 |  | 1.647 |  |
| Cytotoxicity | -2.166 |  | -1.616 |  |
| Response of mononuclear leukocytes | -1.248 |  | -2.531 |  |
| Binding of myeloid cells | -1.255 |  | -2.458 |  |
| Toxicity of cells | -2.315 |  | -1.355 |  |
| Immune response of T lymphocytes | -1.528 |  | -2.1 |  |
| T cell response | -1.278 |  | -2.338 |  |
| Response of lymphocytes | -0.848 |  | -2.368 |  |
| Invasion of malignant tumor | 2.343 |  | 0.829 |  |
| Function of blood cells | 0.458 |  | -2.543 |  |
| Function of leukocytes | 0.629 |  | -2.361 |  |
| Adhesion of endothelial cells | 0.891 |  | -2.059 |  |
| Invasion of tumor | 2.004 |  | 0.924 |  |
| Mitogenesis | 2.794 |  | N/A |  |
| Accumulation of lymphocytes | 0.621 |  | -2.05 |  |
| T cell migration | -2.116 |  | -0.468 |  |
| Cell-mediated response of T lymphocytes | -0.515 |  | -2.028 |  |
| Binding of breast cancer cell lines | 2.166 |  | -0.346 |  |
| Cell movement of T lymphocytes | -2.135 |  | -0.375 |  |
| Response of antigen presenting cells | -0.16 |  | -2.291 |  |
| Cytotoxicity of lymphatic system cells | -2.434 |  | N/A |  |
| Adhesion of blood cells | -0.338 |  | -2.048 |  |
| Cytotoxicity of cells | -2.349 |  | N/A |  |
| Cytotoxicity of lymphocytes | -2.313 |  | N/A |  |
| Binding of endothelial cells | 0.18 |  | -2.128 |  |
| Inflammation of airway | N/A |  | -2.29 |  |
| Migration of vascular cells | 2.028 |  | -0.25 |  |
| Expansion of T lymphocytes | -2.268 |  | N/A |  |
| Adhesion of myeloid cells | 0.061 |  | -2.194 |  |
| Expansion of lymphatic system cells | -2.09 |  | N/A |  |
| MAPKKK cascade | -2.075 |  | N/A |  |
